# Inferring pathways of oxidative folding from pre-folding free energy landscapes of disulfide-rich toxins

**DOI:** 10.1101/2022.10.07.511306

**Authors:** Rachael A. Mansbach, Lara A. Patel, Natalya A. Watson, Jessica Z. Kubicek-Sutherland, S. Gnanakaran

## Abstract

Short, cysteine-rich peptides can exist in stable or metastable structural ensembles due to the number of possible patterns of formation of their disulfide bonds. One interesting subset of this peptide group is the coonotoxins, which are produced by aquatic snails in the family *Conidae*. The *µ* conotoxins, which are antagonists and blockers of the voltage-gated sodium channel, exist in a folding spectrum: on one end of the spectrum are more hirudin-like folders, which form disulfide bonds and then reshuffle them, leading to an ensemble of kinetically trapped isomers–and on the other end are more BPTI-like folders–which form the native disulfide bonds one by one in a particular order, leading to a preponderance of conformations existing in a single stable state. In this article, we employ the composite diffusion map approach to study the unified free energy surface of pre-folding *µ*-conotoxin equilibrium. We identify the two most important nonlinear collective modes of the unified folding landscape and demonstrate that in the absence of their disulfides, the conotoxins can be thought of as largely disordered polymers. A small increase in the number of hydrophobic residues in the protein shifts the free energy landscape towards hydrophobically collapsed coil conformations responsible for cysteine proximity in hirudin-like folders, compared to semi-extended coil conformations with more distal cysteines in BPTI-like folders. Overall, this work sheds important light on the folding processes and free energy landscapes of cysteine-rich peptides and demonstrates the extent to which sequence and length contribute to these landscapes.

## 1 Introduction

Short, cysteine-rich peptide toxins are good candidates for novel therapeutic leads due to their high binding affinity and specificity for different protein receptors involved in cell signaling pathways.^1–7^ A perennial difficulty in rationally designing such peptides for binding to a particular receptor is the difficulty of controlling the resultant fold, since the covalent bonds that form between the disulfides have a strong influence on the structure. Also, the formation of bonds between different cysteine pairs under oxidative folding conditions leads to isomers with multiple patterns of disulfide bonding, which may have different binding specificities and affinities. ^8^

The question of the conformations taken on by short, cysteine-rich peptides is intimately related to the question of protein folding. In particular, there are two folding pathway types proposed for cysteine-rich proteins: “hirudin-like” folding and “BPTI-like” folding. In the former, as research on its namesake protein suggests, disulfide bonds form indiscriminately and later interchange to reach the native state, whereas in the latter, the native disulfide bonds form one at a time, progressing from the unfolded state directly to the native fold.

The current consensus is that the mechanism is sequence and length dependent.^9,10^

We focus on understanding the effects of sequence on the early folding process before any of the disulfide bonds are formed. A high proportion of short, cysteine-rich peptides are predicted to be disordered in the absence of the formed disulfides; however this has not been tested.^11^ Several different techniques have been employed to avoid disulfide reshuffling with concomitant kinetic trapping, including the replacement of disulfide bonds with dicarba bonds or diselenide bonds,^12,13^ the creation of mutants with one or two deleted disulfide bonds that are still stable, ^14,15^ or a combination of these techniques. Here we stress an alternative: that an ensemble of stable or metastable states of different structure significantly expands the possible library of structures for therapeutic design of possible receptor ligands. A thorough understanding of sequence contributions to folding propensity is the first step in using kinetics rather than thermodynamics to control the resultant conformational ensembles. Kinetic control has a rich history of use in materials design applications;^16,17^ in particular it has shown great use in the controlling the assembly of peptides in nanocontrolled metamaterials.^18^–21

In this article (see Fig. 1 for an overview), we use five *µ*-conotoxins as a relevant testing ground to explore whether the free energy landscape of short, disulfide-rich peptides lying closer to the hirudin end of the folding continuum will be significantly more rugged in the absence of disulfide bonds than will the free energy landscape of those that lie closer to the BPTI end. We perform replica exchange molecular dynamics (REMD) with the disulfide bonds disconnected, as has been previously done in the case of several *α* conotoxins^22–24^ to study the effects of solvation in aqueous ionic liquids but not for the purpose of assessing prefolding equilibrium. By employing the diffusion map dimensionality reduction technique ^25–27^ in its composite formulation, ^28^ we are able to directly compare the energy landscapes of the different conotoxins in what is essentially a “shared basis” of the underlying lower-dimensional nonlinear manifold of the configurational space. We find that the free energy landscapes are more similar than we had expected and display a broad, shallow character consistent with disordered polymers, but there is a shift in the populations of the conotoxins corresponding to differences in length, charge patterning, and percentage of hydrophobic residues. We observe that the distance of the energy minimum from the projection of the “native state” in the free energy landscape is predictive of the type of folder. We validate our results via comparison to a previous study on these five *µ* conotoxins that employed largely experimental techniques to order them along the oxidative folding continuum. ^29^

**Figure 1:**
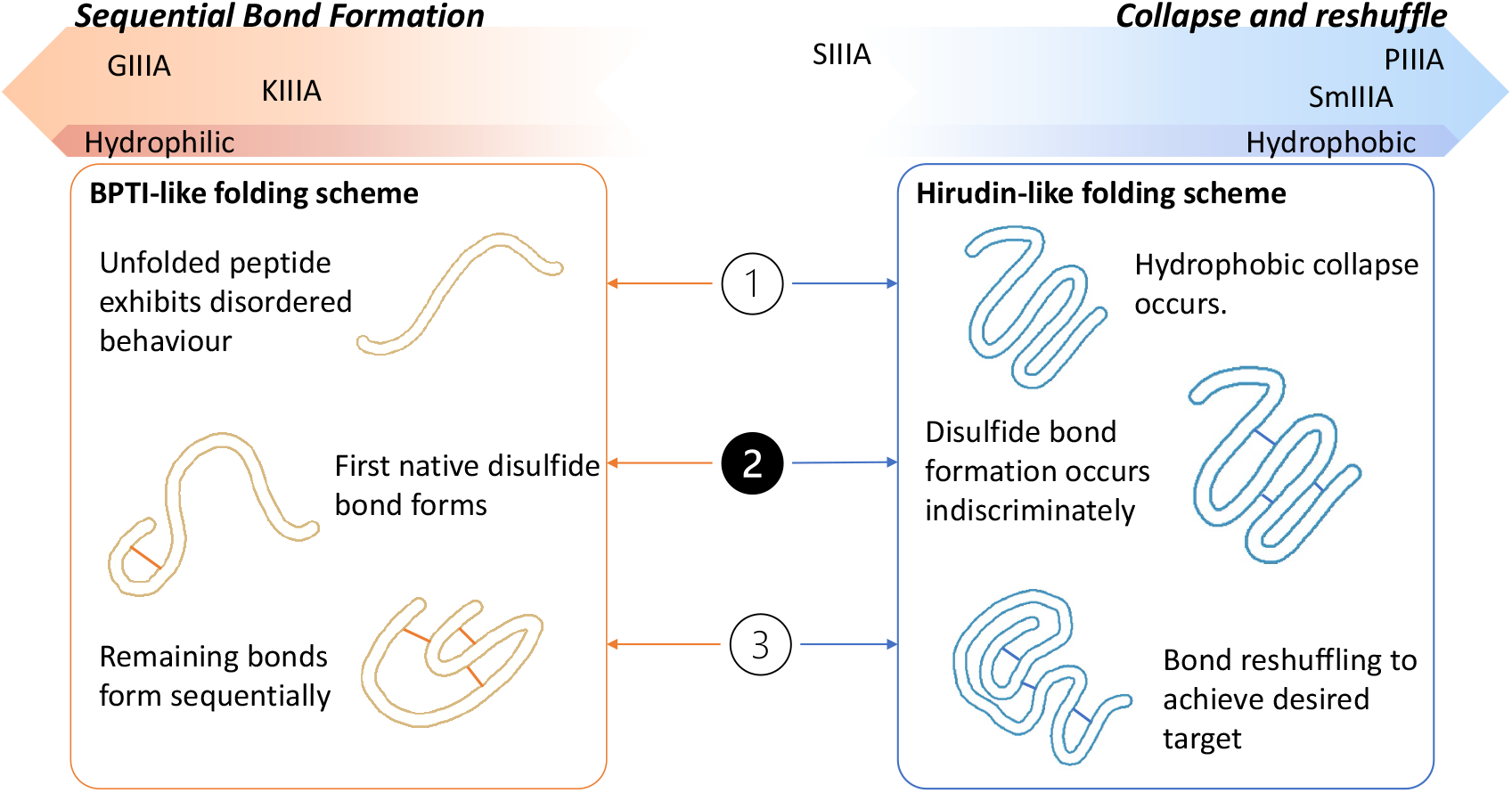
Folding continuum of the conotoxins described in this work. The Sequential Bond Formation (left) scheme follows a mechanism in which the unfolded peptide exists in a relatively disordered state until two cysteine residues come into proximity with each other and form a native bond. Following this, the native bonds are sequentially formed until the peptide is folded. In contrast, the Collapse and Reshuffle (right) scheme follows a mechanism in which the peptide first undergoes hydrophobic collapse, which puts many cysteine residues into proximity, so non-native bonds form quickly, and then undergo reshuffling to achieve the native fold.

## 2 Methods

### 2.1 Composite diffusion maps

Protein configurations may be described as existing in 3*N* -dimensional space, where *N* is the number of atoms in the system. Due to bonded and nonbonded interactions, however, many of the atoms move in strongly collective manners such that typically far fewer than 3*N* variables are required to describe the configurational space of protein and peptide systems. More rigorously, the Mori-Zwanzig formalism states that there exists a lower-dimensional manifold–often in practice a nonlinear one–onto which the 3*N* Cartesian space may be projected without loss of information.^30,31^

Many different algorithms have been developed to project high-dimensional data onto lower dimensional surfaces. Perhaps the simplest is Principal Components Analysis (PCA), in which a spectral decomposition is used on the correlation matrix to identify the (linear) manifolds in order of decreasing variance.^32^ PCA, indeed, has demonstrated great utility in the protein-folding and protein-dynamics world,^33^ but it can suffer from loss of accuracy due to the stringent requirement of projection onto a linear rather than a nonlinear manifold. Algorithms that determine projections onto nonlinear manifolds include Isomaps, locally linear embedding, Laplacian eigenmaps, spectral embedding, multi-dimensional scaling, and others, many of which have been profitably employed for numerous data visualization applications.^34^ More recently, deep learning approaches such as variational autoencoders have also demonstrated their use, particularly with respect to problems of protein folding and dynamics.^35,36^

The diffusion map is a nonlinear dimensionality reduction technique that has the desirable property that, loosely, the distance between points projected into an associated manifold (the “diffusion distance”) corresponds to the inverse of the probability of transitioning between the states they represent in the original high-dimensional space (the “dynamical distance”). This property makes the diffusion map a good technique with which to study dynamical systems, such as proteins, because there is an intuitive interpretation of the description of the system in terms of the modes corresponding to the greatest diffusion-map eigenvalues. ^37,38^ One constructs the diffusion map as follows: ^38^ First, define a distance metric between two datapoints of interest. Prior work in MD simulations has typically employed the backbone root mean square deviation (RMSD) of configuration snapshots that have been translationally and rotationally aligned; however, the only requirement that such a metric must fulfill is that it be a good proxy for *local* dynamical distances. Following this, construct the *R* × *R* pairwise distance matrix **P**, where *R* is the number of datapoints. Next, kernelize and normalize **P** to form a right stochastic Markov matrix,

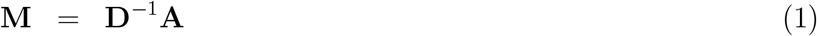

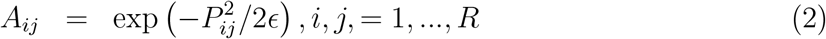

where **D** is the diagonal matrix containing the row sums of **A**,

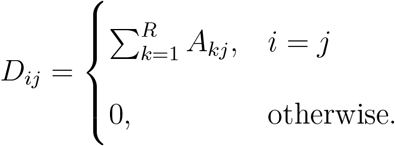

The exponential kernel essentially serves as a soft thresholding that retains only local distances; the variable may be tuned to control the exact size of the local region. The choice of an exponential specifically is of great relevance, since it is the infinitesimal generator of a diffusion process, leading to the interpretation of operator **M** as defining a discrete random walk over the data, meaning that the entries *M*_*ij*_ may be thought of as the probability of transitioning from conformation *i* to conformation *j*.

A spectral decomposition of **M** results in a decreasing series of eigenvalues, *λ*_1_ = 1 ≥ *λ*_2_ ≥ … ≥ *λ*_*R*_, the first of which is trivially equal to 1 by the Markov property of **M**. The corresponding right eigenvectors 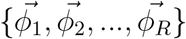 are discrete approximations to the eigenfunctions of the Fokker-Planck operator. ^39^ The first eigenvector is the trivial all-ones vector, because if the **M** matrix is applied often enough the system will go to its stationary probability distribution with the eigenvector of 1 (Markov property). The rest of the eigenvectors correspond to collective motions along the nonlinear manifold of the data. Often there is an identifiable spectral gap after the first few eigenvectors, in which a rapid drop in the eigenvalue spectrum indicates rapidly-changing motions that are effectively slaved to those described by the first few.

We have previously employed a composite diffusion map approach to define an interpretable shared basis for the comparison of free energy surfaces of systems in different environments and with different chemistries and used these to determine the driving forces behind conformational change in a system of alkanes and a system of radially amphiphilic antimicrobial peptides.^28^ The composite approach differs from the traditional diffusion map approach only in that it synthesizes data from systems under different conditions. Free energy surfaces may be constructed in the modes revealed in this way, allowing a direct comparison of the FESs of systems under different conditions. The only requirement is that we must be able to define a sensical distance metric between configurations belonging to systems under different conditions. In addition, once the modes have been identified, new configurations may be embedded in the constructed energy surface by employing the Nystrom Extension procedure.^40,41^

### 2.2 Replica exchange molecular dynamics simulations

Employing Gromacs 5.1.2, ^42^ we conducted replica exchange molecular dynamics (REMD) simulations^43^ of five different *µ* conotoxins at different temperatures, ranging from approximately 280K to 430K (see Table 1 for a more detailed summary), with an exchange rate of approximately 0.3. Peptides were prepared with C terminus amidation and N terminus acetylation, except for peptides terminating in pyroglutamic acid, which used modified parameters for that residue (see Supplemental Information for a more detailed explanation of force field parameters). All disulfide bonds were interactively disconnected using the –ss flag of the `pdb2gmx` Gromacs command. Each conotoxin was solvated in a dodecahedral water box with a principal box vector large enough that it was not expected to self interact (see Table 2 for box vector lengths of the different conotoxins and Sec. A.2 for a description of how these lengths were determined). We used the TIP3P water model.^44^ High energy overlaps in initial configurations were removed by performing steepest descents minimization to remove forces in excess of 1000 kJ/mol. Initial atomic velocities were drawn randomly from a Maxwell distribution. NVT and NPT equilibration runs of 100 ps each were performed, using a velocity rescaling thermostat^45^ and, for the NPT equilibration, a Parrinello-Rahman barostat,^46^ after which temperature, pressure, and density had attained stable values. Production simulations were conducted in the NPT ensemble, employing a Nosé-Hoover thermostat^47^ and a Parrinello-Rahman barostat. Lennard-Jones interactions were smoothly shifted to zero at a cutoff of 1.2 nm. Electrostatic interactions were computed with the Particle Mesh Ewald (PME) method^48^ with a Fourier grid spacing of 0.12 nm and a real-space cutoff of 1.2 nm. We used a leap-frog algorithm^49^ with a 2 fs time step to numerically integrate the equations of motion. Bond-lengths were fixed with the LINCS algorithm.^50^ Conotoxins were modeled with the CHARMM36m force field, ^51^ modified to add parameters for post-translationally-modified residues hydroxyproline and pyroglutamate (see Sec. A.1 for a full discussion of modifications). We employed the CHARMM force field for compatibility with possible future membrane calculations, and specifically employed version CHARMM36m due to its increased ability to model disordered proteins. At the beginning of the production runs and after 500 ns of production, there were no unusual chiralities in the molecules. There is a single cis peptide bond in GIIIA between two hydroxyproline residues, which is unusual, due to the fact that even in hydroxyproline and proline residues, in which cis peptide bonds are less disfavored than in most other amino acid residues, cis peptide bonds are about ten times less likely.^52^ Nonetheless, this cis peptide bond is found in all GIIIA NMR structures, as well as the NMR structures of the closely related GIIIB and GIIIC (PDB IDs 1GIB and 6MJD).^53,54^ Therefore we assume that this unusual bond is due to the sequence and constraints, and we retain it in our models. We do not see any shift to a trans peptide bond over the entire 500 ns of production.

**Table 1:**
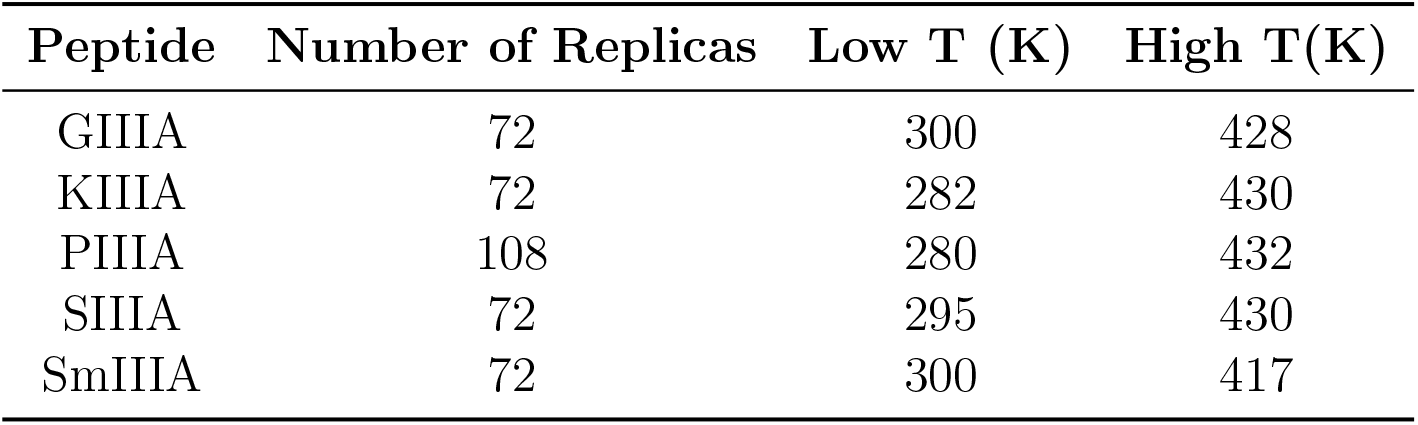
Principal box vector length for dodecahedral boxes for production REMD.

**Table 2:**
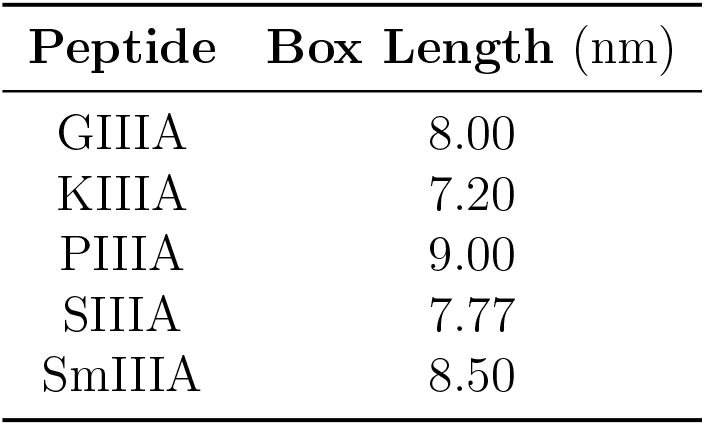
Principal box vector length for dodecahedral boxes for production REMD.

### 2.3 Unbiased molecular dynamics simulations

As part of validating the force field we employed (see Sec. A.1), we also performed unbiased molecular dynamics simulations of the conotoxins with their disulfides connected. Peptides were prepared in the same manner from initial PDB structures but without disconnecting the disulfides. For constrained peptides, which are not expected to deviate strongly from their initial structures, boxes were constructed by padding with 1.2 nm of water around the initial peptide conformers. Production simulations at 300 K were 375 ns in length, of which we discarded the first 75 ns as initial equilibriation.

## 3 Results and discussion

By using the composite diffusion map approach, in the following sections, we identify a two-dimensional unified pre-folding free energy landscape for five *µ* conotoxins: *µ*-SmIIIA, *µ*-KIIIA, *µ*-SIIIA, *µ*-GIIIA, and *µ*-PIIIA (cf. Fig. 1). We show that the two collective variables we identify are related to hydrophobic collapse of short biomolecules as previously observed in a similar manner for alkanes. ^28,38^ We find that all free energy surfaces are quite broad and shallow, indicating disordered polymeric behavior as previously hypothesized, and the lack of persistent secondary structural motifs in most of the toxins supports this. We do find that there are some length, charge, and sequence dependent effects that change the propensity of different peptides for different parts of the landscape, including a persistent *α*-helical motif in one conotoxin (SIIIA) that causes it to depart from predicted behavior. By analyzing the native and non-native cysteine proximities, the proportion of native contacts, and the secondary structural content, we delve more deeply into these effects and demonstrate their relationship to location on the oxidative folding continuum.

### 3.1 Composite diffusion map modes describe folding collapse in two dimensions

#### 3.1.1 Distance metric, kernel size, approximate dimensionality, and functional mode dependency

The first step in constructing the diffusion map is to identify a distance metric that captures small, local perturbations. Previously, for peptides with different sidechain moieties, we employed the RMSD between heavy backbone atoms and considered the differing sidechains to be captured implicitly. ^28^ In this article, we are considering sequences that do not have identical backbones, so we first perform a multiple sequence alignment with Clustal Omega ^55^ and then perform minor adjustments by hand, such that all cysteines are aligned (see Fig. 2)). In addition to the six cysteines, nine residues align with no gaps in the sequence, for a total of 14 aligned residues, which is 14/22 or 63.6% of the longest sequence and 14/16 or 87.5% of the shortest sequence. We define the distance metric of the composite diffusion map to be the RMSD between the C*α* atoms of the nine aligned residues plus the RMSD between all heavy atoms in the cysteine residues, for a total of 45 atoms considered in the RMSD. Although this disproportionately covers the C-terminus side of the sequences, due to the shortness of *µ*-KIIIA, we believe that the focus on aligned cysteine atoms, together with the two cysteine residues no more than three residues away from the N-terminus, provides a good local distance metric. We focus on the cysteine residues due to their importance in forging the necessary disulfide bonds, but we also consider the other nine residues in the alignment to gain a fuller picture of the overall conformations to the extent possible with the sequence differences. Other differences between the sequence are captured implicitly. Although work has been done on capturing solvent rearrangement,^56^ we do not explicitly consider solvent due to computational intractability of constructing a diffusion map representation of the large solvent boxes needed to capture the extended non-folded conformations.

**Figure 2:**
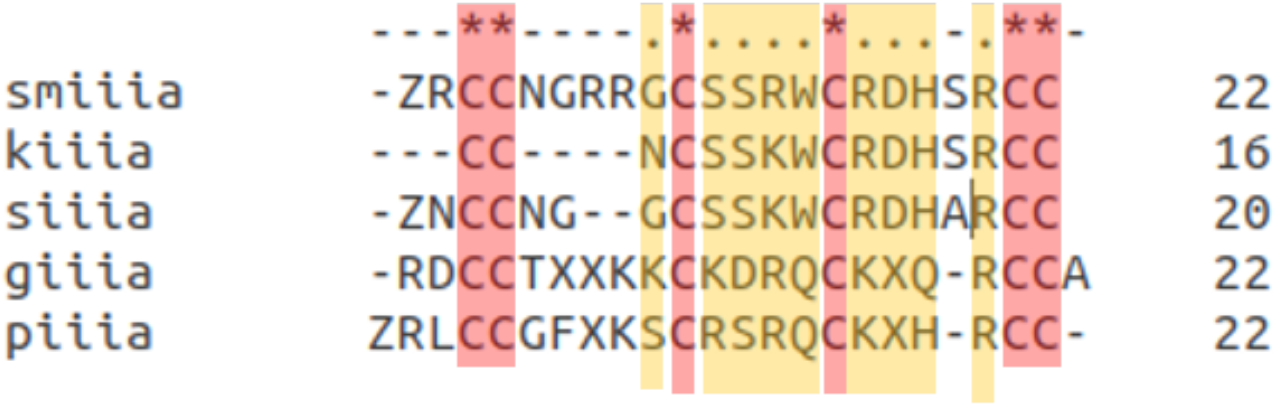
Multiple sequence alignment of the *µ* conotoxin sequences considered in this work. Label on the left; sequences in the center, with dashes representing gaps, and sequence length in the rightmost column. In the top row, dashes represent a column with at least one gap, asterisks represent the six cysteines that are forced to align (columns highlighted in red), and periods represent non-cysteine sequence alignments with no gap (highlighted in yellow). The alignment was performed with Clustal Omega ^55^ and modified slightly by hand to enforce cysteines alignment in all columns.

With the distance metric determined, we proceed to the computation of the diffusion map, which we perform on 10,000 equally-spaced frames from each of the five trajectories for a total of 50,000 data points. The frames were drawn from the second to lowest temperature of each one (301.57, 301.87, 301.90, 301.51, and 301.44 K respectively). Although strictly speaking the simulations were performed at different temperatures, the maximum difference is less than 0.1% and therefore should not cause significant deviations in the constructed diffusion map or free energy surfaces.

We identify an appropriate value of ϵ= 0.1 – the bandwidth of the Gaussian kernel – by following an approach previously described.^28,38,39^ From the log-log plot of ∑*ij A*_*ij*_ versus *ϵ* (see Fig. S6c), we find an approximation for the dimensionality of the system to be 2-3. As expected, *λ*_1_ = 1, corresponding to the trivial eigenvector *ψ*_1_ of the equilibrium distribution. The eigenvalue spectrum (Fig. S6a) shows a spectral gap after *ψ*_4_, the third non-trivial eigenvalue, suggesting a three-dimensional system; however, close inspection of the relationship of the eigenvalues demonstrates a functional dependency between *ψ*_2_ and *ψ*_4_ (Fig. S6b). Such functional dependencies have previously been observed in diffusion map analyses, and we remove this dependence in the same manner as has previously been done,^28,38^ by representing the two variables by their arc-length, *α*_24_, parameterized by fitting to a single second-order polynomial (Fig. S6b, red line). Thus, the overall dimensionality of the system is approximately two, and the system is spanned by two basis vectors, *α*_24_ and *ψ*_3_.

#### 3.1.2 Collective modes correspond to hydrophobic collapse of short polymers

We characterize the two important collective modes of the pre-folding landscape through visualization and identification of pertinent bridge variables (Fig. 3). *α*_24_ is highly correlated with the end to end length (Fig. 3a), with a Pearson correlation coefficient of *ρ* = 0.86, *p <* 0.00001. We were unable to find such a clear bridge variable for *ψ*_3_, but we gain some insight by visually inspecting the relationship between *ψ*_3_ and the relative shape anisotropy, *κ*^2^, defined as,

**Figure 3:**
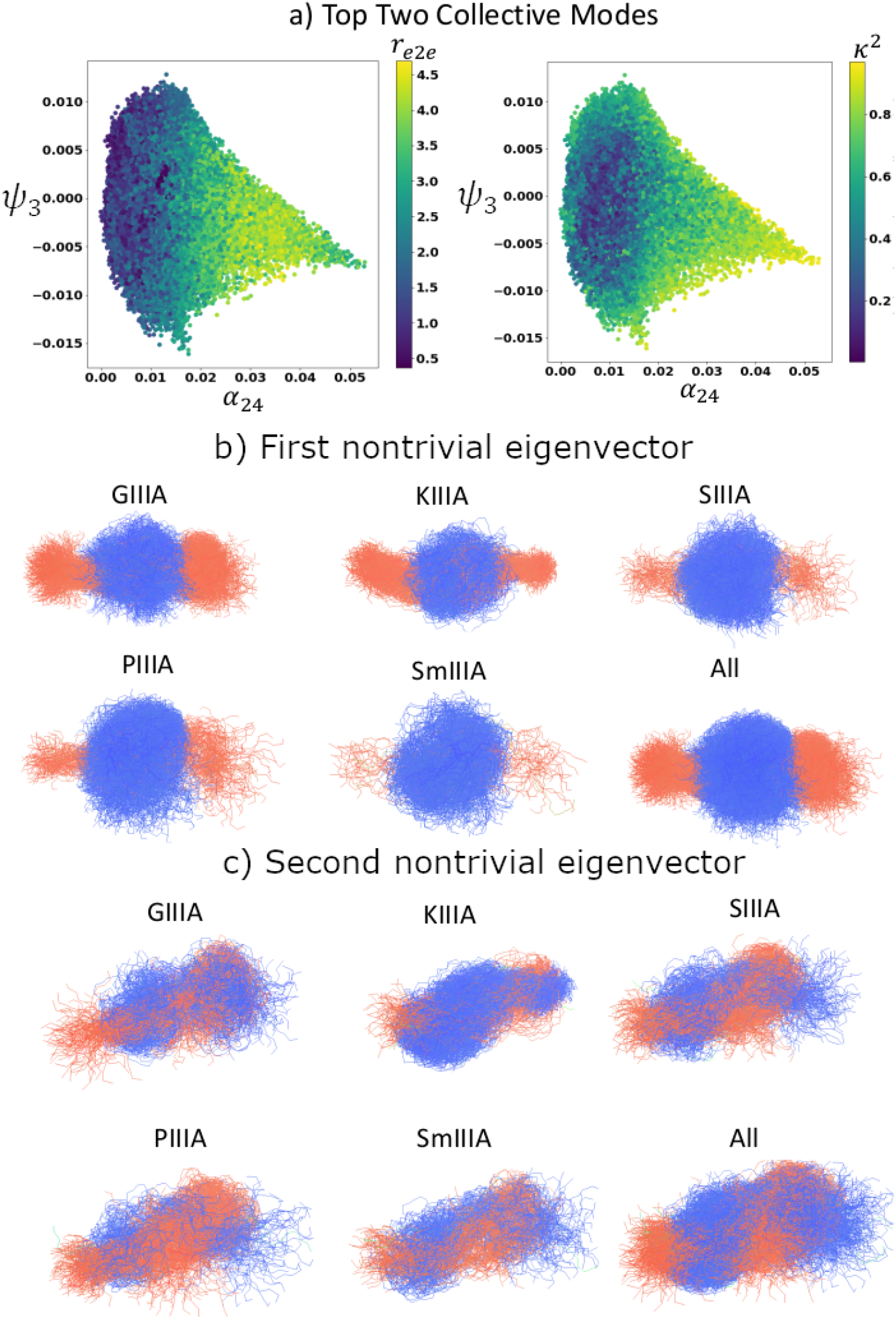
Collective modes identified by the composite diffusion map. In a) we illustrate the configurations projected into the top two collective modes colored by *r*_*e*2*e*_, end to end length, (left panel) and *κ*^2^, relative shape anisotropy (right panel). In b)-c), we visualize the two modes through demonstrating the configurations that contribute most strongly to each one. Red corresponds to configurations contributing strongly to one extremum of the mode (low *ψ* value) while blue corresponds to configurations contributing strongly to the other extremum (high *ψ* value).

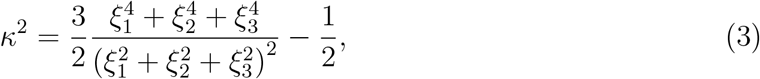

where *ξ*_*i*_ is the *i*th principal moment of the gyration tensor. The relative shape anisotropy is, roughly speaking, a measure of how spherical or linear a conformation is. It runs from 0 −meaning that the object demonstrates tetrahedral or higher order symmetry (sphere-like) − to 1, meaning the object is linear.^57^

Thus, we can see that the configurations are most anisotropic when *α*_24_ is maximal, corresponding to highly extended configurations, and that *ψ*_3_ covers its greatest range of values for small values of *α*_24_, i.e. collapsed configurations demonstrate a greater range of *ψ*_3_ values. In addition, *ψ*_3_ is approximately symmetric in *κ*^2^, and the greatest isotropy is observed for small values of *ψ*_3_.

In Fig. 3b-c, we gain further insight through visualizing the top two eigenvectors. (The third eigenvector is visualized in Fig. S7b; it is correlated with the top eigenvector, as mentioned.) We utilize a method we developed previously, based on a similar one developed for visualization of diffusion map modes in image processing.^28,58^ In brief, a properly-weighted sum of the configurations projected into the mode may be thought of as the single “most” representative configuration. We may visualize such a sum in configurational space–where the single-number representation is not informative–by coloring configurations in accordance with their weights in the sum. We thus demonstrate the largest contributions to the most representative configuration to gain an intuitive understanding of the mode even in the absence of a bridge variable or linear mapping. Because this visualization was derived in terms of the eigenvectors, we visualize *ψ*_2_ and *ψ*_4_ separately, rather than *α*_24_. For ease of visualization, we display the top 10% of configurations in all five systems that correspond to the modes of interest.

From Fig. 3b, we observe that the first collective mode is indeed related to the end-to-end length, for the configurations contributing high mode values (red, right end of *α*_24_) are highly extended, almost linear conformations and those contributing low mode values (blue, left end of *α*_24_) are highly collapsed, almost spherical conformations on the other. Since we visualize the top 10% of all configurations, not per peptide, the lack of extended configurations of SIIIA and SmIIIA demonstrates that they occupy a less extended space overall than GIIIA or KIIIA (as does PIIIA, to a lesser extent.)

From Fig. 3c, we observe that the second collective mode (*ψ*_3_) constitutes a transition from a kink on one end of the peptide to a kink on the other end of the peptide. We note a weak anti-correlation of the absolute value of this mode with molecular acylindricity (with a Pearson correlation coefficient of *ρ* = -0.19, *p <* 0.0001). We may rationalize this as follows: when the mode has a large magnitude, it is more highly kinked and thus more cylindrical, while when it has a small magnitude, it is closer to spherical collapse. This also explains the observations with relation to *κ*^2^, because configurations with small contributions to the mode are relatively U-shaped and isotropic but configurations with large contributions (i.e. at the extremes of the modes) are anisotropic with a “hook” on the right or left end of the peptide. A similar collective mode has previously been observed in alkanes solvated in water: ^28^ it is related to hydrophobic collapse through demonstrating a zipper-like mechanism that allows a biomolecule to expel water molecules from its center as it collapses. We do note, however, that the different peptides demonstrate different propensities to sample one side of the hook over the other, with SmIIIA in particular contributing comparatively less to this mode, perhaps because it spends more of its time populating the spherical collapsed configurations identified in the previous figure. To gain further insight into the differential populations of the configurations, we construct free energy surfaces in the identified collective modes.

### 3.2 Shallow free energy surfaces indicate disorder

We constructed two-dimensional free energy surfaces in the collective modes with the `plot_free_energy` function of the PyEmma “plots” package^59^ using ten levels and twenty bins. We constructed one-dimensional free energy surfaces separately fitting the principal modes, *α*_24_ and *ψ*_3_, with a Gaussian kernel density estimation G using the `gaussian_kde` function from the scipy.stats Python package^60^ to approximate the density of points at each place on the surface for a direct numerical comparison at the same points. From this approximation, we construct the free energy surface as,

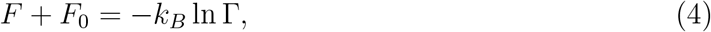

where we threshold all values of G Γ *<*= 10^*-*6^ to zero, and *F*_0_ is an arbitrary constant of the free energy.

In Fig. 4, we illustrate the two-dimensional free energy surfaces of the five studied *µ*-conotoxins in the shared collective basis identified by the composite diffusion map, and in Fig. S8, we illustrate the corresponding one-dimensional fits. The full free energy surface spans about 5*k*_*B*_*T*, and the differences in occupancy are relatively small and largely within the scale of thermal fluctuations. All free energy surfaces display relatively flat, wide basins, which suggests that the peptides are quite disordered, since they do not demonstrate a strong preference for one configuration over another. This is supported by a secondary structure analysis (cf. Sec. 3.4) that demonstrates that most of the peptides are primarily in a random coil state.

**Figure 4:**
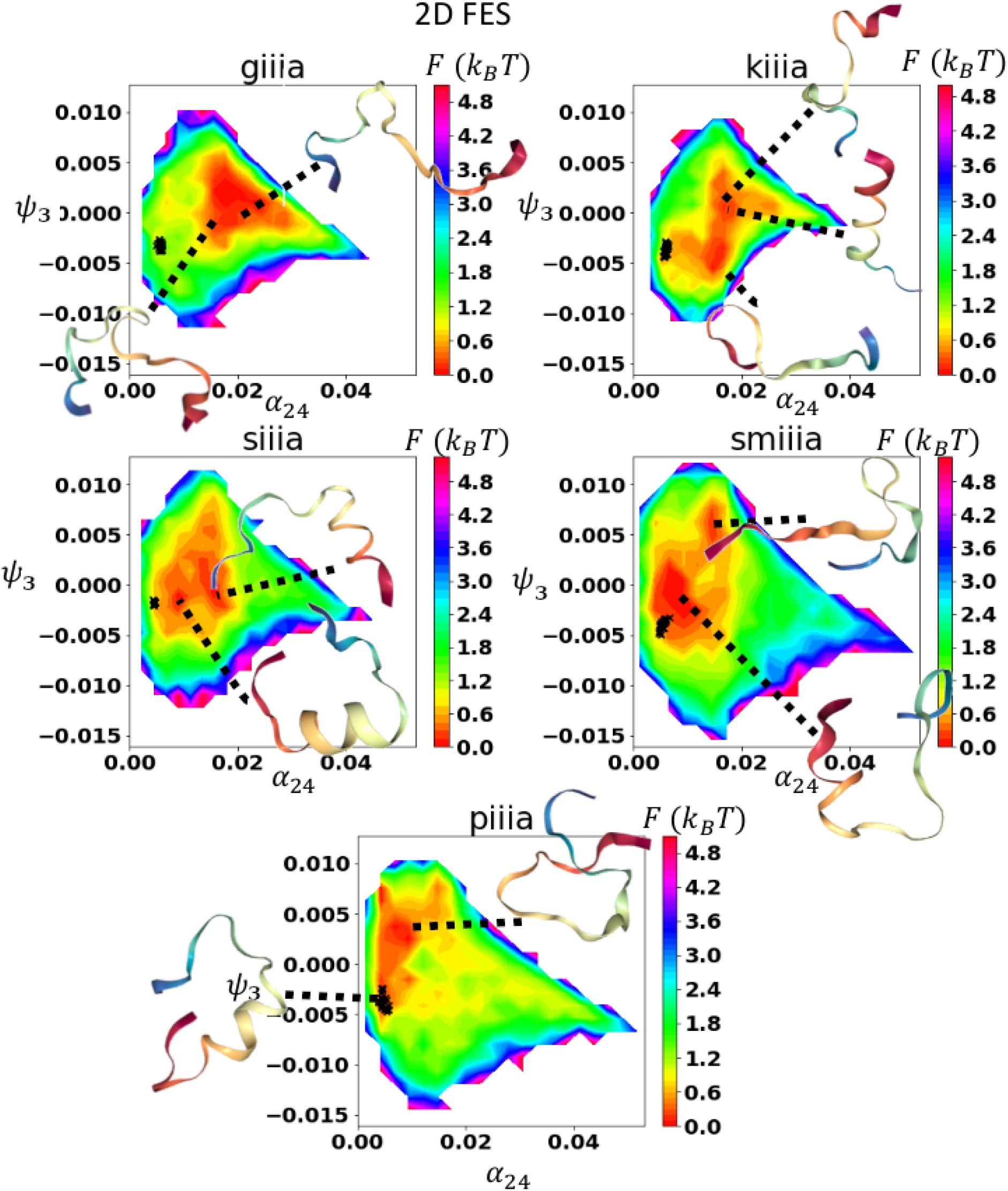
We present the two-dimensional free energy surfaces projected into the two top collective modes of the system using the Pyemma package.^59^ Black Xs demonstrate the location of representative frames from disulfide-connected simulations in the native state, projected into the FES using the Nystrom extension. ^40,41^ Representative conformations produced with NGLview. ^61^ We demonstrate two KIIIA conformations from one basin to show that there is some helical content but the basin is not solely comprised of helical conformers. Note that PIIIA’s secondary minimum is occupied by the native state conformers.

The two peptides that were previously identified as the most hirudin-like, PIIIA and SmIIIA, demonstrate a higher propensity for collapsed conformations (Table 3), with both the average location of the free energy minima and their approximate ranges shifted to lower values of *α*_24_ compared to GIIIA and KIIIA. They also demonstrate a shift towards lower *ψ*_3_ values in comparison to the other three, corresponding to a tendency to populate only one end of the hook-slide mode. In addition, GIIIA (the least hirudin-like) and KIIIA (the shortest) peptides tend to sample a smaller proportion of *ψ*_3_: in Fig. 4 GIIIA and KIIIA show overall less extended basins about their minima and in Fig. S8, their free energy surfaces are significantly higher at the extreme ends. Thus, although indicative of disorder, the peptide pre-folding free energy surfaces do demonstrate different propensities indicative of peptide folding type: more hirudin-like peptides are on-average more collapsed and also explore a greater portion of the total free energy landscape. Furthermore, these differences clearly arise from sequence (or charge) and length-based considerations, since the disulfides are not yet connected.

**Table 3:**
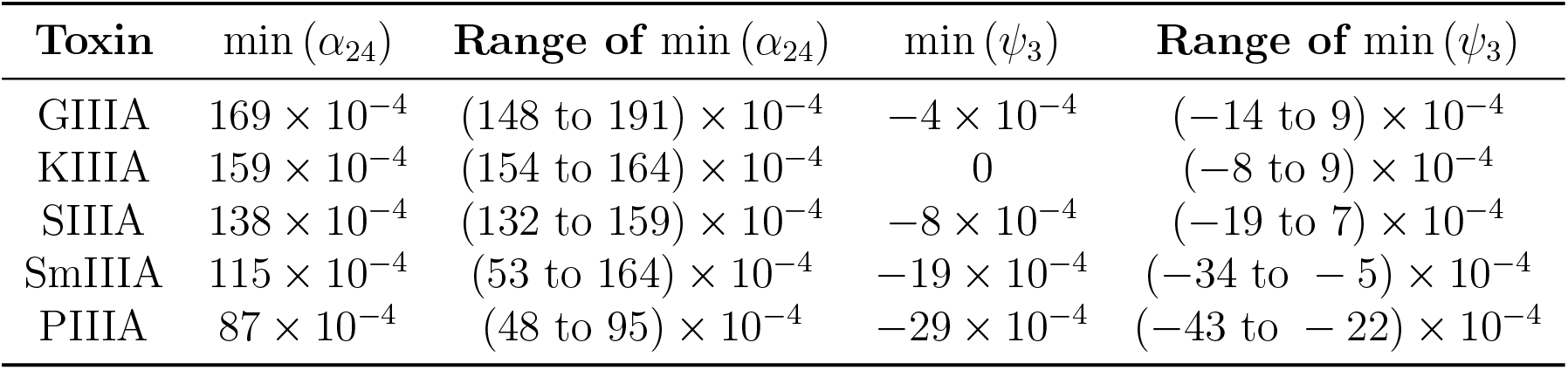
Locations of minima for the free energy surfaces of the two dominant collective modes, *α*_24_ and *ψ*_3_. Since the minima were collected by fitting a Gaussian KDE to a set of numerical values, we report significant figures out to the step size between the points 5× 10^*-*4^ for *α*_24_ and 3× 10^*-*4^ for *ψ*_3_. We also report the range of values for which the free energy lies within error bars of the minimum, where the error bars are computed by block averaging with *N* = 5.

The approximate “distance” between the locations (described in Table 3) of the free energy minima and the locations (described in Table 4) of the projections of observed “native” states (simulations performed with disulfides connected from NMR structures from the PDB) corresponds roughly with the expected ranking from hirudin-like to BPTI-like (cf. Fig. 5). These distances are computed in the top two modes of the diffusion map; they are so-called “diffusion distances,” which correspond to dynamical distances in the original high-dimensional space^39^ and thus can be thought of as how much a conformation lying in the free energy minimum of pre-folding must evolve in order to reach the native state. We observe that GIIIA – the most BPTI-like folder – has a minimum furthest from its native state, with KIIIA, SIIIA, SmIIIA, and PIIIA closer in descending order, though when the ranges are taken into account the differentiation is closer to a binary with GIIIA, KIIIA, and SIIIA in the “far” group and SmIIIA and PIIIA in the “near” one. It should be also noted that SmIIIA in particular has a very large range of free energy minimum values. Breaking the distances down into *α*_24_ distances in one dimension and *ψ*_3_ distances in another dimension (Fig. 5b), we observe that the downward trend is somewhat clearer for *α*_24_, whereas the binary trend is more evident for *ψ*_3_. In other words, GIIIA generally remains more extended with its disulfides unconnected than it is in the native state, with KIIIA second, then SIIIA, SmIIIA, and PIIIA. In the hook-slide mechanism, GIIIA, KIIIA, and SIIIA are about the same distance away (demonstrating greater isotropy in the disconnected state than in the native state), while PIIIA and SmIIIA are about the same distance away, but they are not very far away at all (demonstrating a similar level of isotropy with disulfides connected and disconnected). Overall, the more hirudin-like folders exist close to their native states even with the disulfides disconnected; the more BPTI-like a peptide is, the more its occupation of the energy surface shifts away. The interpretation of this is quite straightforward: hirudin-like folders collapse, connect disulfides indiscriminately, and reshuffle, whereas BPTI-like folders do not collapse until the first disulfide connects.

**Figure 5:**
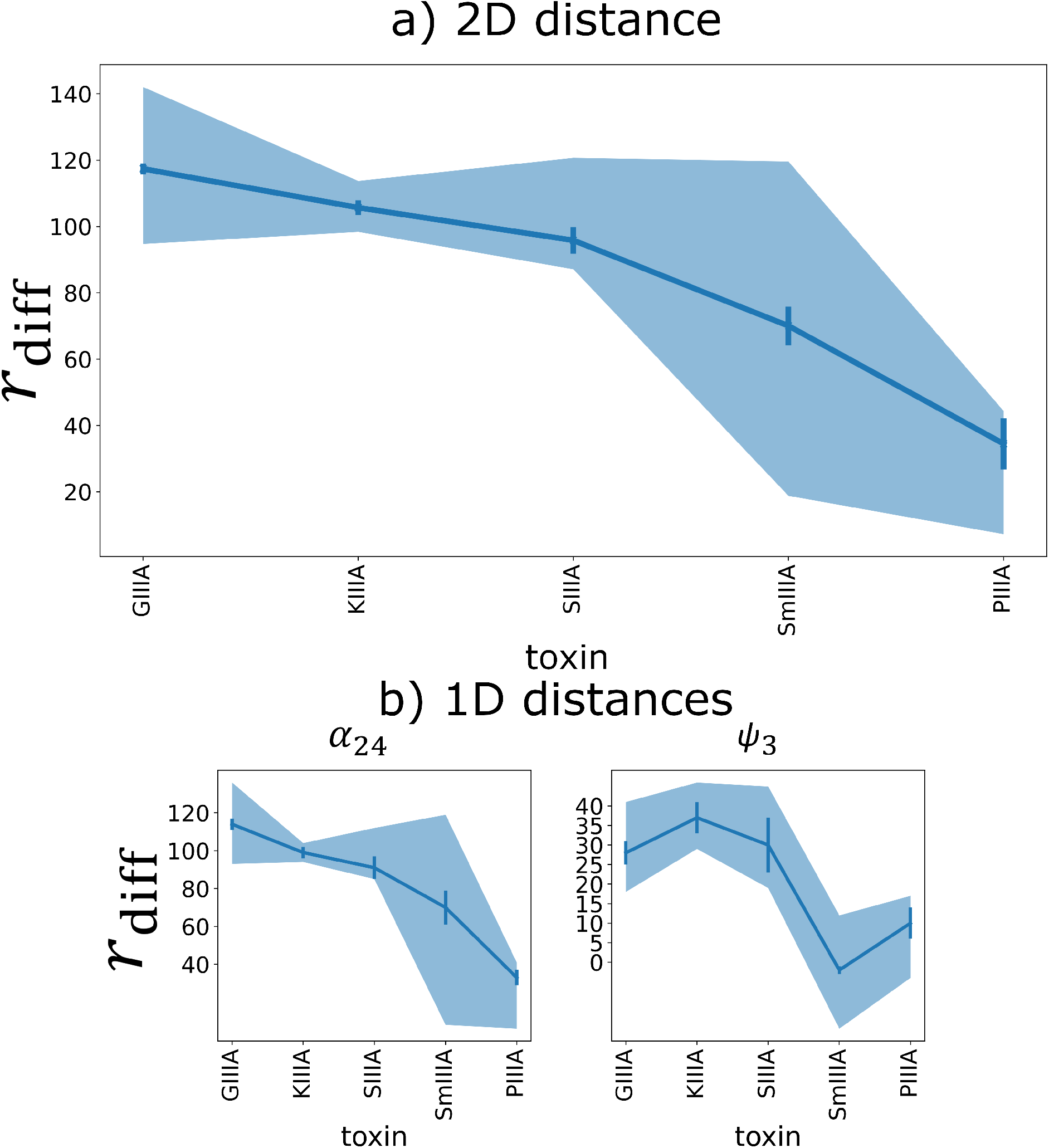
Diffusion distances *r*_diff_ between free energy minima and average native-state projection into diffusion map coordinates. Each point is calculated as the (a) two-dimensional and (b) one-dimensional distance between the approximate location of the minima identified from the one-dimensional FES fits and the average value of the projected native states calculated from the Nystrom extension, both calculated in terms of the top two eigenmodes of the diffusion map. Error bars represent standard error propagation of the standard deviations reported in Table 4, while the shaded area is calculated from the distance between the lower and upper ends of the ranges shown in Table 3.

**Table 4:**
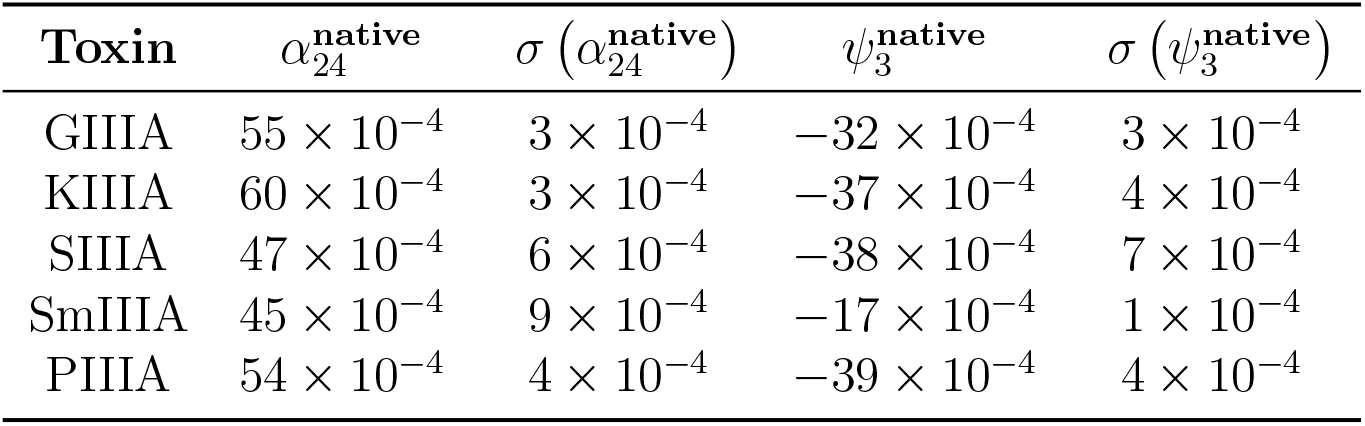
Average locations of “native state” projected with the Nystrom extension ^40,41^ onto the free energy surfaces of the two dominant collective modes, *α*_24_ and *ψ*_3_. We report the mean and standard deviation of 30 subsampled snapshots.

### 3.3 Contacts and Cysteine Proximity Indicate Differences Between Hirudin-like and BPTI-like Folders

To further assess what information we can glean from the pre-folding surfaces, we assess the proximity of cysteine residues and proportion of contacts formed in different parts of the unified free energy landscape.

In Fig. 6, we show as black triangles projected onto the free energy surface, configurations where one native cysteine contact is transiently “formed”, that is where the two sulfur atoms of the cysteine residues involved in one native disulfide are within a cutoff distance of 5 angstroms. Sulfur proximity has previously been used to estimate propensities of different disulfide isomers in AuIB.^62^ We show as cyan triangles configurations where two native cysteine contacts are “formed.” We do not observe any configurations where all three native contacts are formed.

**Figure 6:**
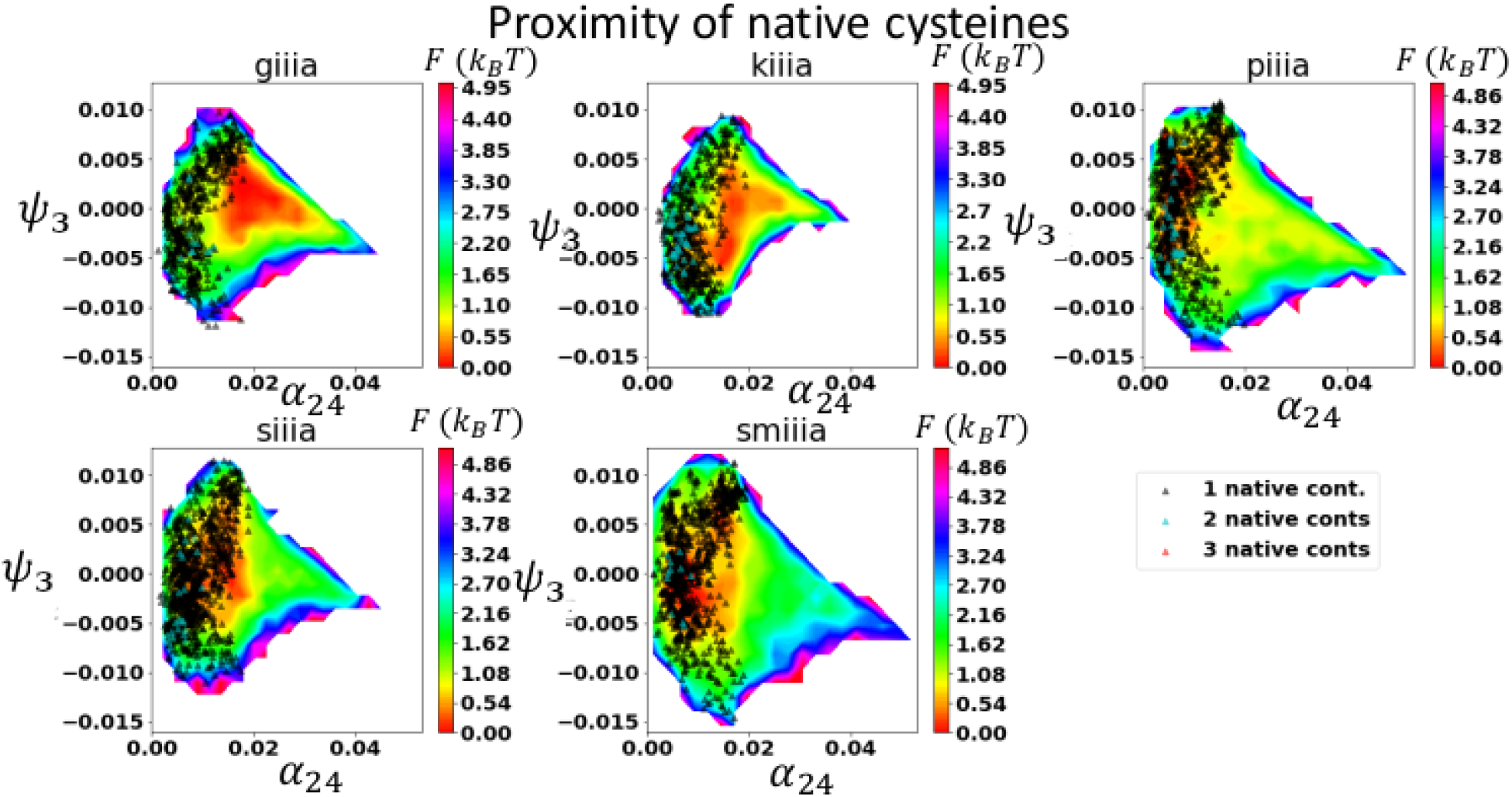
Differential population of transient native bonds in free energy landscapes. In this figure, we indicate the portions of the free energy surfaces where cysteines are in close proximity (within a cutoff of 5 angstroms) and thus bonds might in principle be able to form between them. Black triangles indicate close proximity between one set of cysteines; and cyan between two.

Observations of the free energy surfaces demonstrate that the minima of GIIIA and KIIIA overlap little with the proximity of any native cysteine contacts, SIIIA displays moderate overlap (its secondary minimum), and PIIIA and SmIIIA demonstrate strong overlap. The minima of all five peptides overlap with the proximity of many single non-native cysteine contacts (Fig. S9), but the minima of GIIIA and KIIIA do not overlap with the proximity of three non-native cysteines and do not overlap much with the proximity of two. We note that the minima of PIIIA and SmIIIA do correspond with the proximity of three non-native cysteines, and while the largest minimum basin of SIIIA does not, there is a secondary small local minima basin that does. (The proximity of one non-native cysteine takes place across a large portion of the energy landscape, presumably due to the short lengths of the peptides in question and the fact that the cysteines are not necessarily far from one another in the sequence.)

Based on these observations, we conclude that PIIIA and SmIIIA do indeed undergo hirudin-like collapse and it is evident from their free-energy surfaces, since their minima overlap with the proximity and thus the likely formation of many non-native cysteine bonds, whereas GIIIA and KIIIA undergo more BPTI-like collapse, where formation of single non-native bonds is possible initially but they do not collapse further in the absence of formation of the native bonds to facilitate folding. It appears that SIIIA may undergo both forms of collapse, with a small subpopulation collapsing and reshuffling disulfides and a larger population driven by the formation of the native disulfides.

We also consider contact formation in general, not limited to cysteine proximity (Fig. 7, Fig. S10, Fig. S11). As expected, there are greater numbers of contacts formed when the peptides have collapsed (Fig. 7a); however, the fraction of total formed contacts that are native is on-average higher in the extended conformations (Fig. 7b), which confirms that collapse tends to be largely indiscriminate; heightened numbers of native contacts in the collapsed conformations are largely due solely to heightened proximity rather than structural formation in the absence of disulfides.

**Figure 7:**
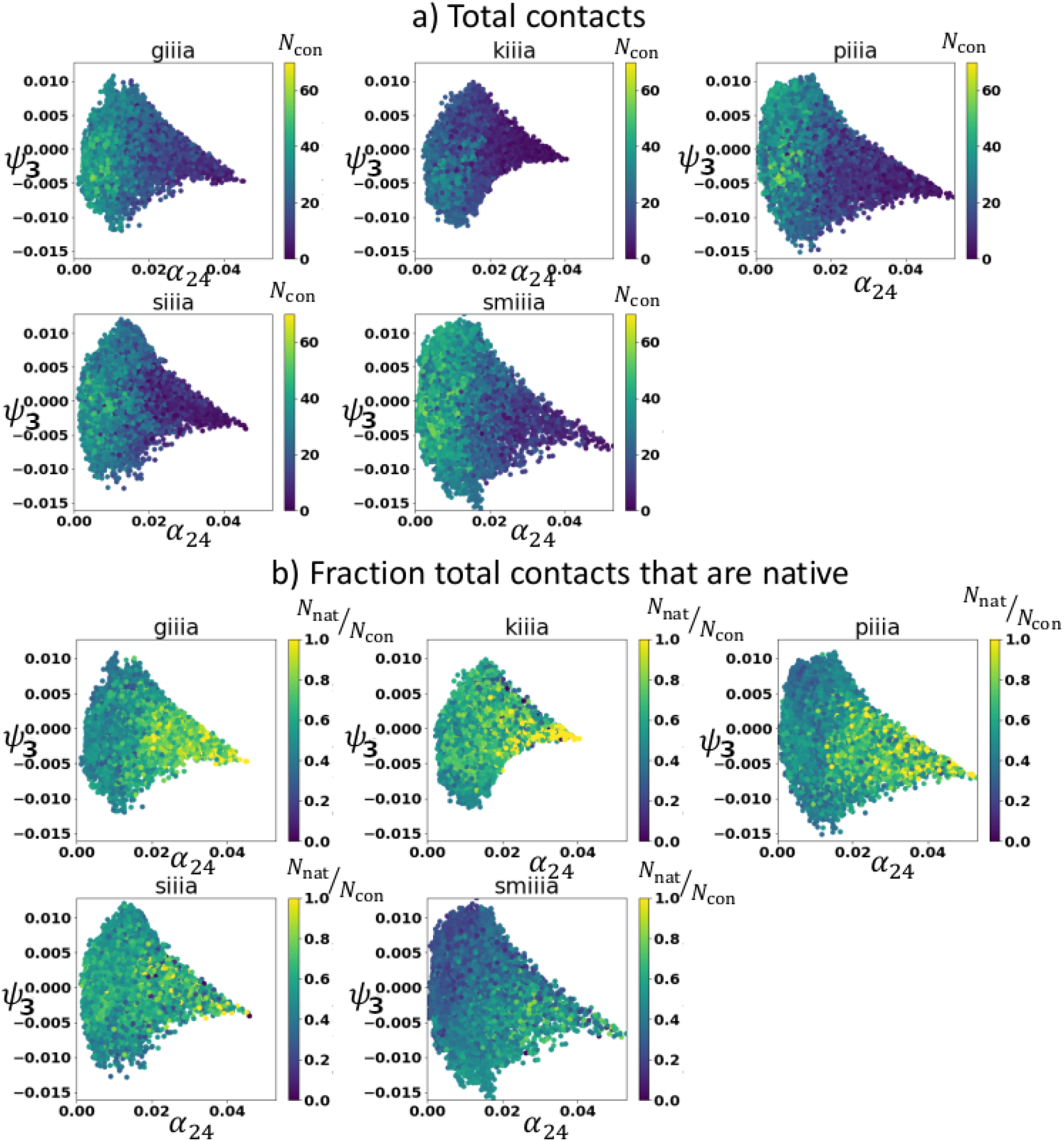
Total contacts. In this plot, we color the two top collective modes by (a) the total number of contacts and (b) the fraction of those total contacts that are native. We observe that in general the areas with lower numbers of total contacts have a higher fraction that are native, corresponding to more extended states.

### 3.4 Secondary structure propensity differences between BPTI-like and hirudin-like peptides: the odd case of SIIIA

Previous work on the folding landscapes of disulfide-rich proteins has indicated that one of the primary factors playing a role in the type of folding collapse that occurs is the relative structural stability of the subdomains of the protein, ^9^ with greater stability corresponding to more BPTI-like, sequential folding. To assess whether this holds true for short peptides and to further investigate the extent of disorder in the conotoxins, we analyze the secondary structural content of the conformations in different parts of the unified free energy landscape.

In Fig. 8, we illustrate the average probability of secondary structure formation per residue over the course of the simulations. We first note that there is a higher percentage of random coil for GIIIA, KIIIA, and PIIIA (∼ 60%) than for SIIIA or SmIIIA (∼ 40%). For most of the proteins, however, the structure that is observed is largely simple coil-like bends and turns, with SIIIA demonstrating a moderately persistent (20% probability) helical motif near its C-terminus, corresponding to a metastable state close to its native one (cf. Fig. 4). Of this secondary structure, only the helical motif in SIIIA and the bend at the fifth and sixth residues of GIIIA represent significant motifs matching the NMR structures (cf. the NMR structure at the top of each plot and Fig. S12 in the Supplementary Information.)

**Figure 8:**
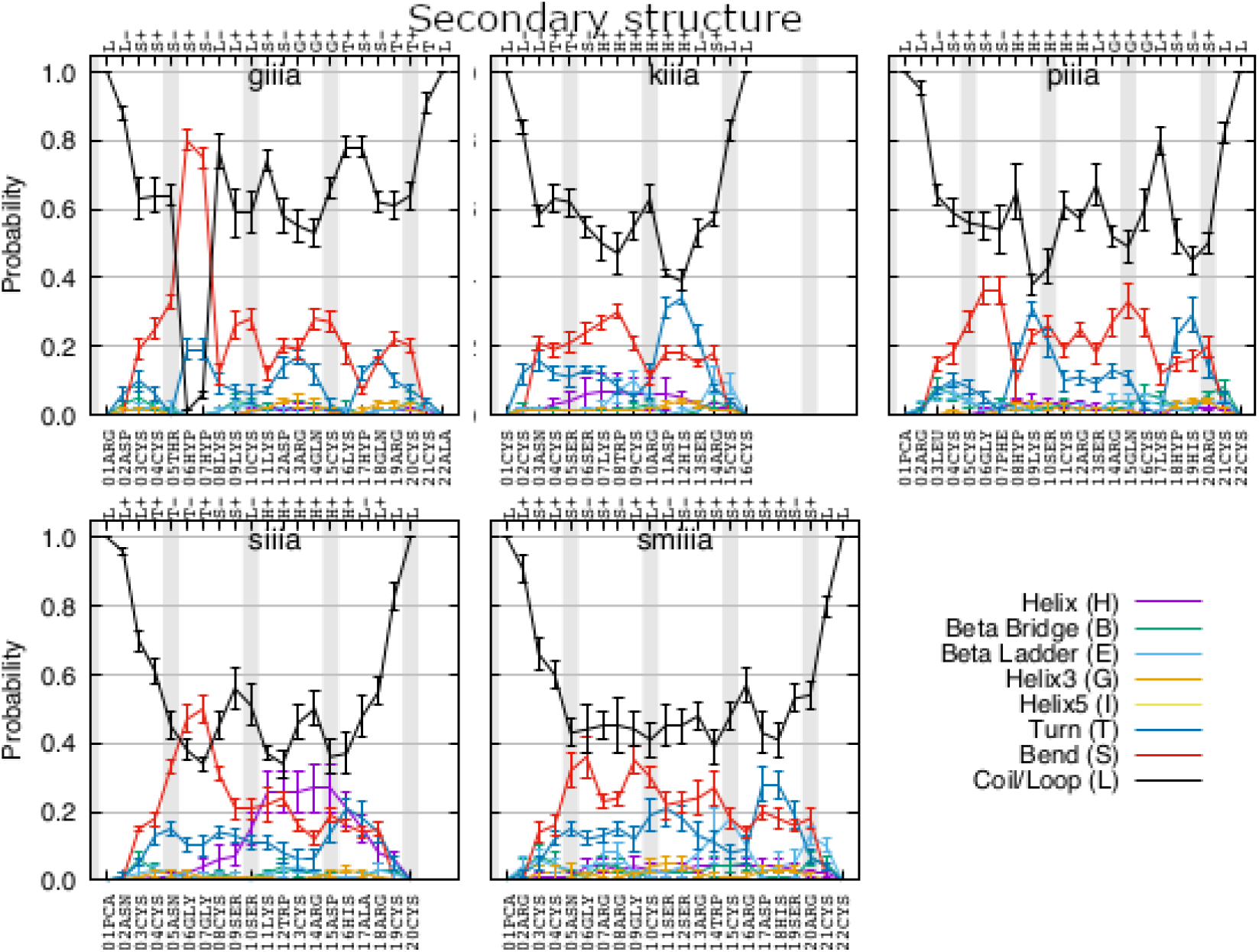
Secondary structure propensities of the *µ*-conotoxins. Average secondary structure per residue. Secondary structure of reference NMR structure shown above the corresponding residues in each respective plot.

Taken together, this tells us that KIIIA maintains a disordered but extended structure, GIIIA maintains a semi-ordered and extended structure, SIIIA splits into subpopulations of extended disordered and collapsed semi-ordered, and PIIIA and SmIIIA form disordered collapsed structures.

### 3.5 Sequence analysis demonstrates the importance of hydrophobic residues

Previous work has suggested that the large number of basic residues in SmIIIA and PIIIA contributes to their tendency to form multiple isomers, but no explanation has been suggested for why GIIIA in contrast does not, and the literature on the matter is somewhat contradictory.^29^ Now that we have conclusively demonstrated that the conotoxins are semi- or fully disordered in the absence of the connecting disulfides, we assess their properties from the perspective of intrinsically disordered proteins (IDPs), whose properties may be predicted based largely on the charge patterning in the sequence.^63^ We calculate the fraction of positive and negative charges in the sequence and plot the five sequences on an IDP phase diagram (see Fig. 9). Based on charge considerations alone, we can see that all the conotoxins are correctly predicted to have some polyampholytic coil behavior, and that PIIIA and SIIIA are both predicted (correctly) to display some globule-like behavior. GIIIA is correctly predicted on this basis to display less globule-like behavior; however SmIIIA is predicted to display less globule-like behavior than KIIIA, so although this assessment is qualitatively accurate in terms of the overall conotoxin behaviors, it is not predictive of the specific folding types.

**Figure 9:**
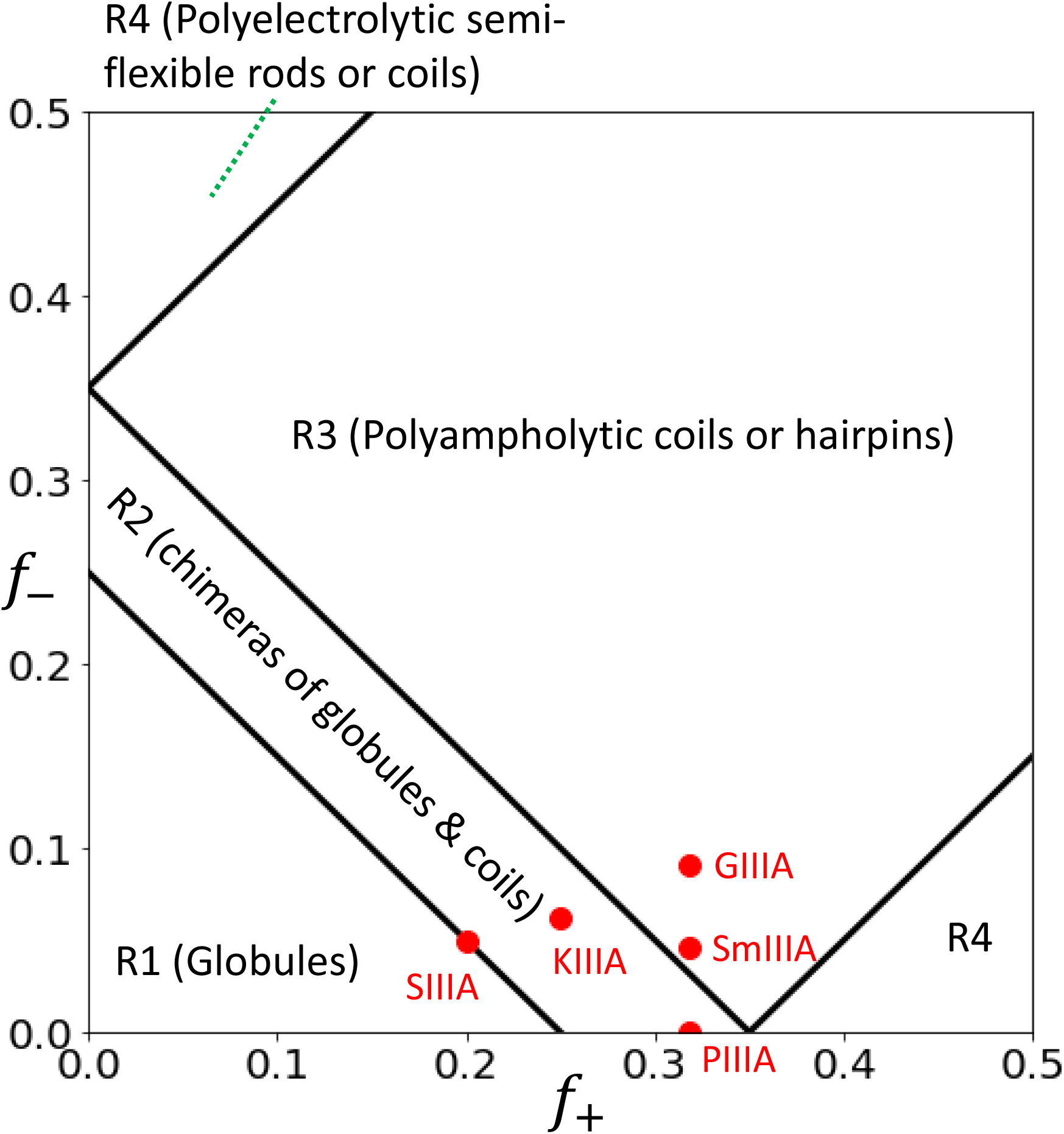
Sequence charge analysis. We plot the fraction of negative charges in each sequence versus the fraction of positive charges in each sequence for the conotoxins (labeled red points) and show where they fall within predefined regions of an approximate phase diagram for intrinsically disordered proteins.^63^

If, however, we also consider the fraction of hydrophobic residues in each peptide, we find that SmIIIA contains about 14%, SIIIA 25%, and PIIIA 18%, in comparison to KIIIA (6%) and GIIIA (0%). This suggests that SmIIIA, PIIIA, and SIIIA have at least some tendency to populate more collapsed regions of the free energy landscape because they can essentially “wrap” around a modest hydrophobic core, a driving force lacking in KIIIA and GIIIA. Overall, our analysis demonstrates that a preponderance of the observed conotoxin properties, including folding type, can be explained by appealing to a simple charge and hydrophobicity analysis. SIIIA, with its double-well potential and demonstrated characteristics of both folding types, is harder to predict in this manner and may be better described in a less coarse-grained way.

## 4 Conclusions

Conotoxins, like many toxins, are rich in cysteines, which allows the possibility of forming different patterns of connectivity during oxidative folding. These different disulfide connectivities can have a drastic impact on the structure and function of conotoxins. Since disulfide bonds have a dominant influence on peptide conformations, it is likely that disulfide isomers may expose different binding interfaces to their target receptors. For the *µ*-conotoxins considered in this study, which have six cysteines, 15 different disulfide isomers are possible, although many of them are not observed experimentally, particularly for BPTI-like folders. An even more important point is the question of whether the dominant disulfide connectivity is the same *in vivo* and *in vitro*. A limited number of studies in the literature have suggested that multiple isomers are possible under both conditions. It is, however, difficult to gain insight on the folding of conotoxins *in vivo* as a number of complicating factors come into play in comparison with the situation *in vitro*. Folding, which occurs inside the venom gland, is influenced by the oxidative conditions, chaperons, crowding, accessibility of disulfide isomerase enzymes, and the peptides’ own signaling or precursor motifs.

In this study, we seek to infer the disulfide connectivities from the energy landscapes of the linear peptides representing the sequences of five *µ* conotoxins. We have performed REMD simulations of the conotoxins in a reduced environment with their disulfide bonds unconnected. Using the composite diffusion map technique, we have identified the primary collective modes, identified their thermodynamic implications, and constructed the free energy surfaces of the conotoxins within a unified pre-folding equilibrium landscape. We demonstrate that the landscape is that of disordered biomolecules, with part of the landscape related to simple hydrophobic collapse, as expected of hirudin-like folders. Conotoxins predicted to be hirudin-like folders disproportionately populate the collapsed part of the landscape. Differences in length and in sequence correspond to small shifts in the population within the landscape. In addition, we are largely able to recapitulate experimental results on ranking of peptides in terms of BPTI-like versus hirudin-like collapse and show that a good predictor of disordered collapse in short, disulfide-rich peptides is distance of the free energy minimum from the projected location of the native state in a unified free energy landscape.

We may understand the biophysical underpinnings of these landscapes through considering the amino acid properties of the sequences, both in a general way and in a more detailed way. In a more general way, we have showed that all the conotoxins are predicted based on charge content to be disordered polymers with some coil-like characteristics and that most of the observed behavior can be explained by assessing the percentage of hydrophobic residues within the sequence – conotoxins with a non-negligible percentage of hydrophobic residues are hirudin-like folders expressing hydrophobic collapse around a modest hydrophobic core, whereas those in which hydrophobic residues are largely absent are BPTI-like folders that display extended character in solution and presumably more significant collapse upon the connection of their first disulfide bond.

The notable exception to this pattern is SIIIA, which experimentally and based on the location of its primary free energy minimum would be a BPTI-like folder, but based on its hydrophobicity and charge patterning would be more hirudin-like. It does display a secondary minimum that lies closer to the predicted location of a hirudin-like folder, and we propose that this double-well character may make the experimental results more difficult to interpret. It is also possible that the semi-persistent helical motif in the half of the protein near its C terminus leads to departure from predicted behavior, since it makes SIIIA the most structured of the toxins and the least likely to be well-described from an IDP perspective. In addition, we note the beginnings of formation of that terminal helix in the stable, more extended part of its free energy landscape (cf. the sample conformation in Fig. 4) suggesting that it does help stabilize SIIIA, leading to more BPTI-like behavior through stability of a subdomain, rather than through the lack of a hydrophobic core, as in KIIIA and GIIIA. Overall, our results partially confirm that the relative structural stability of the subdomains of a protein correspond to BPTI-like folding, but we demonstrate that this is a sufficient rather than a necessary condition.

The *µ*-conotoxin PIIIA has been one of the most well-characterized in terms of its disulfide isomers. A recent study employed a combination of 2D-NMR and MS/MS to characterize the 15 different possible isomers Heimer et al. ^64^. NMR ensembles confirmed that most isomers, including the native state, demonstrated a compact and relatively rigid fold despite their differences in structure.^64^ This compactness is consistent with our findings that PIIIA exhibits the most hirudin-like folding that happens after the collapse of the peptide towards a compact ensemble. The highly compact native isomer is considered to be the most active one against the Nav 1.4 channel; however, as noted, experimentally it is remarkably difficult to clearly separate isomeric mixtures to make a conclusive statement.

Finally, it is worth asking whether the simulations considered here reveal any insights on the general question of how much influence the sequence between cysteines plays on the disulfide connectivities. As has been shown in conotoxins with two disulfide bonds, the chosen solvent and temperature would certainly influence the folding a specific isomer. ^65^

The current simulations instead consider the much simpler case of aqueous solvent assisted folding of the linear peptides of five *µ*-conotoxins. Regardless, it still provides some insights that can serve as hypotheses for future studies. We hypothesize that the peptide sequence can play a critical role in BPTI-like folders such as SIIIA. The peptide sequence may not be as critical, other than for driving the entire peptide towards the compact structure, in hirudin-like folders such as PIIIA. It is still possible that proximal amino acids next to cysteines may play a role in scrambling of disulfides. Consequently, with hirudin-like folders, all three disulfide bonds may be necessary for it to be functionally active, requiring some sort of scrambling of disulfides till the right connectivity is attained. A study that used both experiments and computations, showed that in PIIIA, the three disulfide bonds are required to produce the effective inhibition of Nav1.4. Interestingly, removal of any one of the disulfides reduced the affinity.^66^ In contrast, BPTI-like folders may not require all three disulfides as they undergo sequential disulfide formation. One could speculate that an isomer missing the last disulfide bond may still maintain a reasonable conformation that may be closer to the native conformation. This appears to be the case with KIIIA, which can still be functional with just two disulfide bonds.^67^

Although we have presented this paper in the context of what is known of oxidative folding of disulfide-rich peptides and peptide toxins, many open questions remain, particularly in terms of the disulfide connectivities. One particularly important point is the continued challenge posed by the experimental detection of different isomers. Conventional analytical techniques such as high-performance liquid chromatography (HPLC), nuclear magnetic resonance (NMR) and circular dichroism (CD) may not be adequate. It is not clear how effective the ELISA based on an Epitope-Specific Monoclonal Antibody can be on distinguishing the isomers. Only recently, Diana Imhof and colleagues demonstrated the technical feasibility of LC-TIMS-MS towards identification of individual isomers. They were able to distinguish several 2- and 3-disulfide- bonded isomers of PIIIA.^68^

This article lays the groundwork for further free-energy-based analyses of short cysteine-rich toxins. One limitation is the use of the diffusion map, which does not explicitly return a mapping from the conformational space to the primary collective modes and requires the use of ad hoc measures such as the Nystrom extension to project new configurations into the learned subspace. To address this limitation, in future we plan to parameterize a variational autoencoder that will allow for more natural extension to new peptide sequences. Our demonstration of the similarities of the free energy landscapes, however, show that it should be possible in the future to predict landscapes for similar toxins with far less computational effort. Furthermore, we have shown a clear throughline from the pre-folding landscape to the full folding landscape, which may be achieved through parameterizing a reactive force field specific to disulfide-rich toxins or potentially through the use of emerging deep learning techniques.

Overall, this work sheds important light on the molecular-level determinants of the folding of disulfide-rich peptides and represents a step towards a unified predictor for the development of novel toxin-based therapeutics that rely on a given disulfide bonding pattern.

## Acknowledgement

RAM acknowledges a Los Alamos Director’s Postdoctoral Fellowship. This research used resources provided by the Los Alamos National Laboratory Institutional Computing Program, which is supported by the U.S. Department of Energy National Nuclear Security Administration under Contract No. 89233218CNA000001. This research was undertaken, in part, thanks to funding from the Canada Research Chairs Program. The authors thank Dr. Chris Neale, Dr. Cesar Lopez, and Dr. Amy Migliori for fruitful discussions.

## Supporting Information Available

Modified force field details and verification, and thirteen supplementary figures are available. This material is available free of charge via the Internet at http://pubs.acs.org/.

## Notes

### Competing Interest Statement

The authors have declared no competing interest.

