## Supporting Information for "Inferring pathways of oxidative folding from pre-folding free energy landscapes of disulfide-rich toxins"

### A REMD Details

#### A.1 Force field modification

Due to the incorporation into the studied peptides of post-translationally-modified residues hydroxyproline and pyroglutamic acid, it was necessary to add parameters to the CHARMM36m force field to parameterize them. Initial parameters were generated by employing CGenFF version 1.0.0, force field version 3.0.1[69,70] on the pyroglutamate residue connected to its neighboring arginine and on the hydroxyproline alone. The force field files, containing the modified residues, have been provided as part of the Supporting Information. Partial charges on pyroglutamate were retained from CGenFF, with the charge on the C atom modified from 0.54e to 0.499e to neutralize it, a change of only 7.5%, which results in a charge that is more similar to the charge in the CHARMM36m glutamate residue, which is likely to possess a similar charge distribution. Due to its strong similarity to proline (differing only by the replacement of a single hydrogen with a hydroxyl group), hydroxyproline was represented by analogy using the proline parameters supplemented with the CGenFF parameters and partial charges for the carbon connected to the hydroxyl and its other hydrogen. An extra charge of -0.039e was placed on the oxygen of the hydroxyl compared to the CGenFF parameters in order to neutralize it, a change of only 6%. Bond and dihedral parameters containing at least one atom using CGenFF assigned types were assigned from the CGenFF parameters. Angle parameters containing at least two atoms using CGenFF assigned types were assigned from the CGenFF parameters.

##### A.1.1 Validation of modified force field

In order to assure that parameters were reasonable, we ran 120 ns simulations (see below for detailed description) of GIIIA at 450 K and compared the bond, angle, and dihedral distributions containing all identical atoms of that with a modified peptide we label GIIIA-pro, identical except that all three hydroxyprolines were replaced with standard proline residues. The bond and angle distributions are not significantly different: using a 2-sample KS test, we do not reject the null hypothesis that they were drawn from the same distribution at a significance of 0.05). The dihedral distributions are reasonably similar: although we do reject the null hypothesis for some, visual inspection of the others shows high similarity, and it is a reasonable assumption that the dihedrals are long-range enough to show some differences due to the residues being non-identical.

We also ran 120 ns simulations of the other $\mu$ conotoxins, and in Fig. S1, we show a Ramachandran plot of the backbone dihedrals of all five $\mu$ conotoxins with disconnected disulfides and observe that as we would expect they lie primarily in the marginally allowed and allowed regions of the plot. We observe that the space sampled is very similar for all five toxins, suggesting that they adopt similar conformations in the absence of their disulfide bonds.

In addition to testing the bonded parameters, we also ran 375 ns simulations of all peptides with the disulfide bonds connected at 298 K (see below for detailed description) and assessed their conformations through a Ramachandran analysis and an assessment of the conformational deviation from NMR structures. All analyses were performed on the final 300 ns of simulation.

In Fig. S2, we show the average root-mean-square-deviation (RMSD) versus time for each of the different $\mu$ conotoxins from their corresponding NMR conformers. Overall, $\mu$-KIIIA demonstrates the least deviation, with an average RMSD of $\langle r_{\mathrm{RMSD}} \rangle = 1.014 \pm 0.001$ Å, then $\mu$-SmIIIA, with an average RMSD of $\langle r_{\mathrm{RMSD}} \rangle = 2.277 \pm 0.002$ Å, then $\mu$-GIIIA, with an average RMSD of $\langle r_{\mathrm{RMSD}} \rangle = 2.239 \pm 0.003$ Å, $\mu$-PIIIA, with an average RMSD of $\langle r_{\mathrm{RMSD}} \rangle = 3.228 \pm 0.003$, and finally $\mu$-SIIIA, with an average RMSD of $\langle r_{\mathrm{RMSD}} \rangle = 3.348 \pm 0.001$ Å. The values we derive are roughly similar to those observed for similar systems in Paul George et al.[73] (GIIIA 24% larger, KIIIA 7% smaller, PIIIA 40% larger, SIIIA 30% larger, and

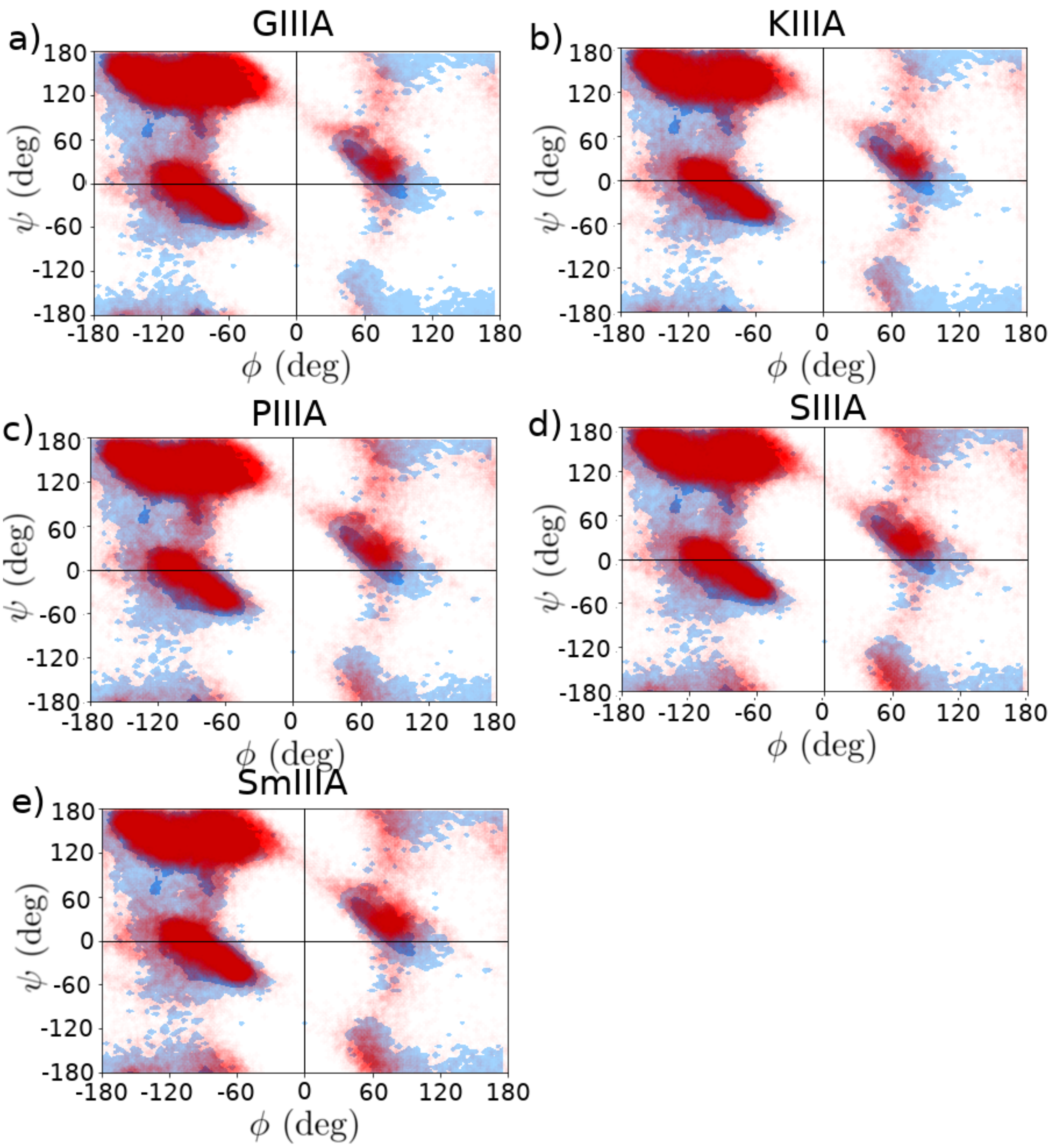


Figure S1: Ramachandran plots of backbone dihedrals for $\mu$ conotoxin (a) GIIIA, (b) KIIIA, (c) PIIIA, (d) SIIIA, and (e) SmIIIA, taken from 60 ns of MD simulations at 450 K with unconstrained dihedrals (red). In dark blue are the "allowed" regions and in light blue the "marginally allowed" regions. Images rendered using the `Ramachandran` function in the MDAnalysis python package v0.19.2.[71,72]

SmIIIA 16% smaller), although a different force field was used, the measurement in that case performed over a third of the time, and our deviation measured with respect to the backbone as opposed to the alpha carbons. This relatively good agreement suggests that our reparameterized force field well captures the expected behavior of the $\mu$ conotoxins. We also note a relatively good agreement in Ramachandran angles sampled in the simulations with those seen in the NMR conformers (Fig. S3). Although there is a larger sampled space for the MD simulations, this is to be expected, as they explore a larger portion of the space, and we note that they remain largely in the allowed or marginally allowed regions of the Ramachandran plot. The one exception is an unusual concentration of ($\phi$,$\psi$) angles in the approximate region running from (0,-60) to (0,60) for $\mu$-SIIIA (Fig. S3d), which visual inspection of the simulation reveals to be due to an unusual backbone configuration of the LYS11 residue, apparently due to the strain of the knotted disulfides on the system. SIIIA contains only a single modified residue, and it is the pyroglutamic acid at the termini, which does not strongly interact with LYS11. Visual comparison between selected NMR structures and final snapshots from MD simulations (Fig. S4d) shows that SIIIA in particular demonstrates a loop movement and helix disruption in the area of the LYS11 residue, leading to greater structural disruption in comparison to the other four.

### A.2 Choice of box sizes

For each conotoxin, we conducted initial standard MD simulations of 120 ns at 450 K, starting from extended conformations produced by employing end-to-end pulling simulations for 500 ps at a rate of 10 nm/ns, with a harmonic pulling force constant of $k$ = 1000 kJ-mol$^{-1}$-nm$^{-2}$. Each conotoxin was placed in a cubic water box of a size larger than twice the LJ cutoff for these simulations (1.2 nm) plus the longest observed distance between two atoms in the extended conformation as measured in VMD[81] (see Table S1). Parameters for the simulations were the same as those used for the REMD simulations described in Sec. 2.2. From the final 60 ns of each simulation, we computed the approximate largest distance

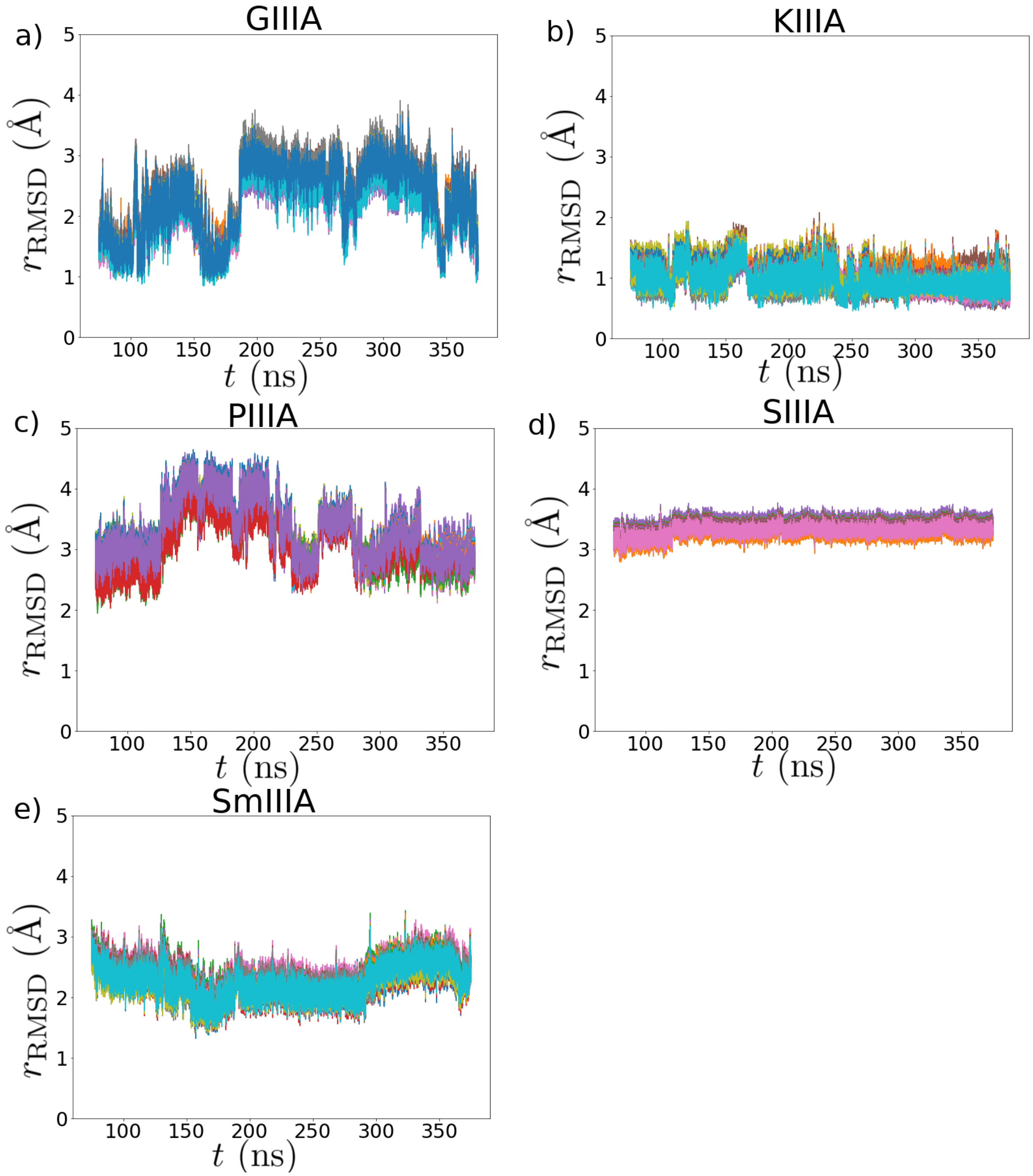


Figure S2: Average root-mean-square-deviation (RMSD) in angstroms from NMR structures versus time in ns of unbiased MD simulations at 298 K with disulfide bonds connected for $\mu$ conotoxin (a) GIIIA, (b) KIIIA, (c) PIIIA, (d) SIIIA, and (e) SmIIIA. Different colors identify different NMR structures. We employed the NMR structures taken from (a) PDB ID 1TCG and 1TCJ [74] (b) PDB ID 2LXG (c) structure ID S00159 [75] on the Conoserver, [76] (d) BMRB structure 20025 [77] and (e) PDB ID 1Q2J. [78] We use the same structures as Paul George et al. [73] for a more direct comparison. Different colors denote different NMR conformers.

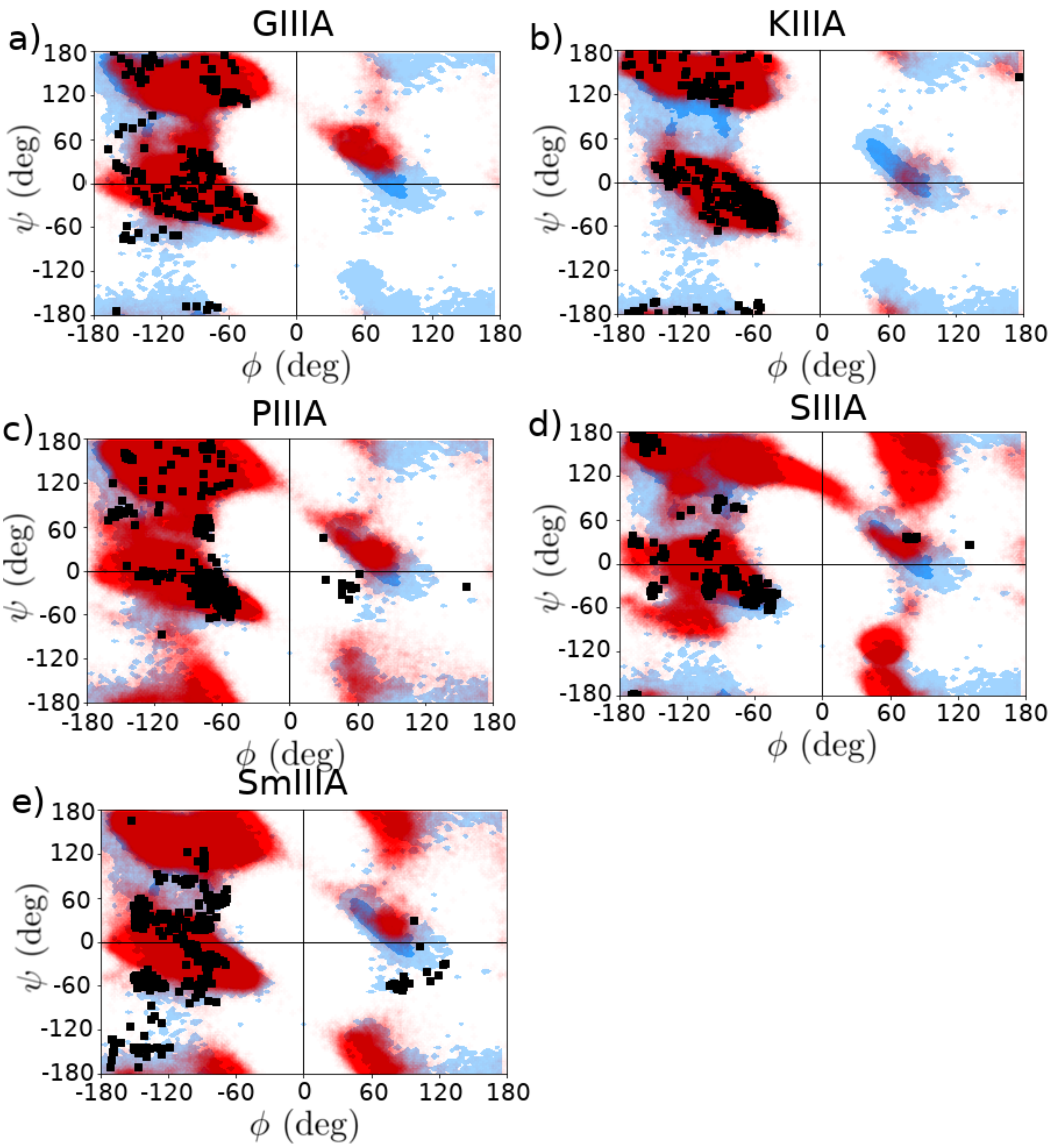


Figure S3: Ramachandran plots of backbone dihedrals for $\mu$ conotoxin (a) GIIIA, (b) KIIIA, (c) PIIIA, (d) SIIIA, and (e) SmIIIA, comparing NMR structures (black squares) with 300 ns MD simulations with connected disulfides (red). In dark blue are the "allowed" regions and in light blue the "marginally allowed" regions. Images rendered using the `Ramachandran` function in the MDAnalysis python package v0.19.2.[71,72]

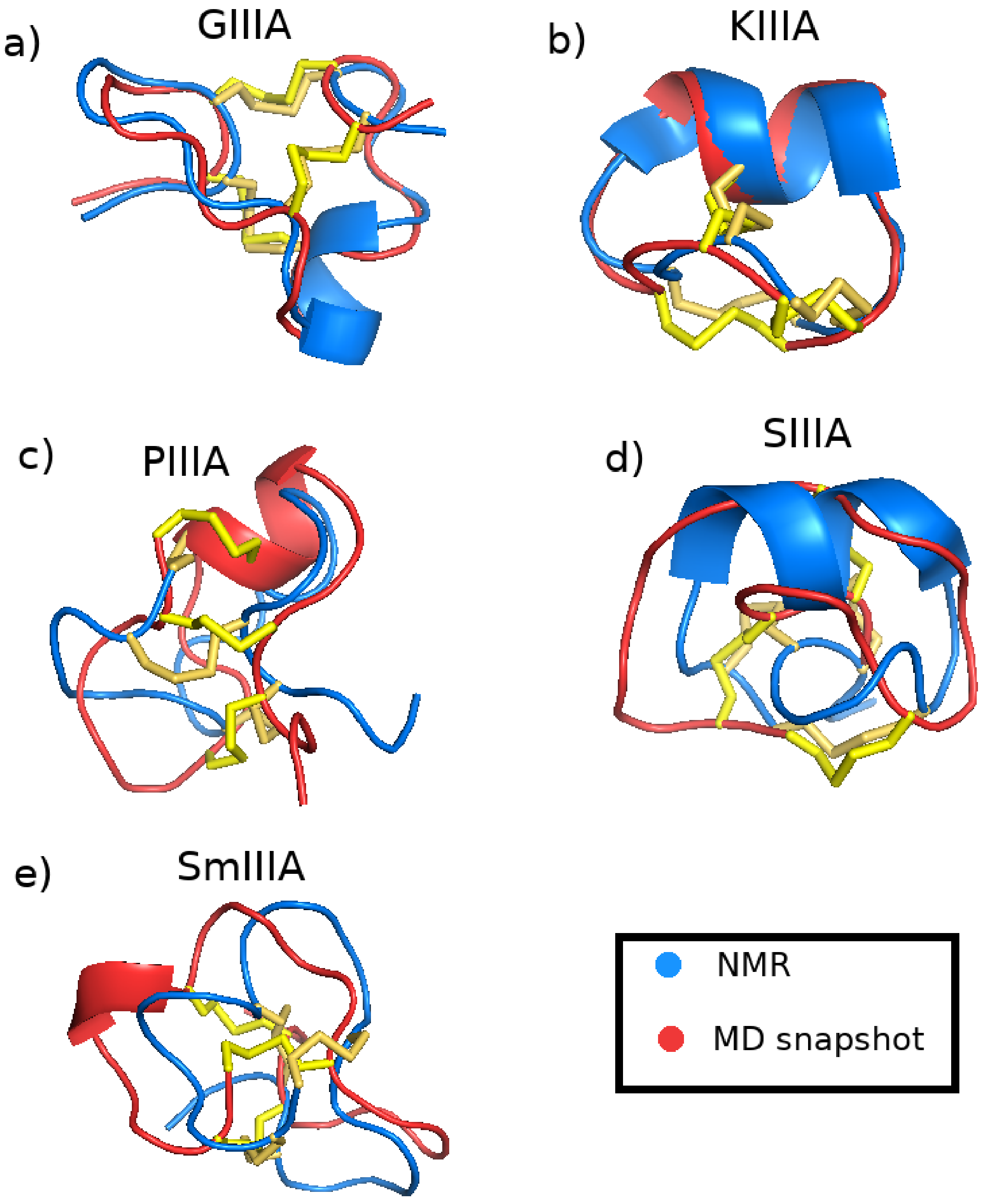


Figure S4: Comparison of selected NMR configurations to molecular dynamics snapshots taken after 375 ns of simulation at 298 K for $\mu$ conotoxin (a) GIIIA, (b) KIIIA, (c) PIIIA, (d) SIIIA, and (e) SmIIIA. NMR structures rendered in blue, MD snapshots rendered in red, with disulfide connections shown as yellow sticks. All images rendered with PyMol.[79]

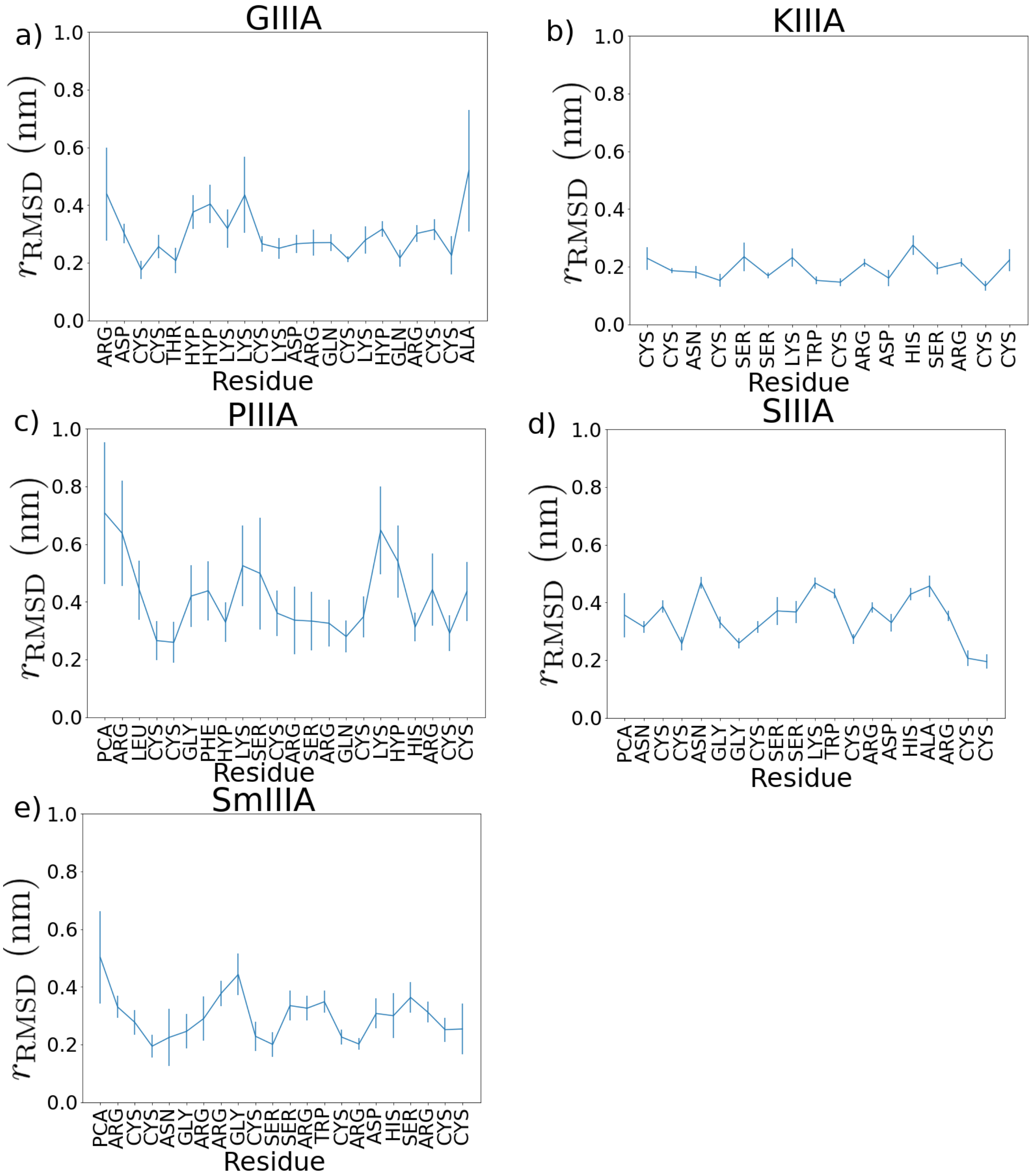


Figure S5: Average root-mean-square-deviation (RMSD) in nm per backbone residue from initial configuration for $\mu$ conotoxin (a) GIIIA, (b) KIIIA, (c) PIIIA, (d) SIIIA, and (e) SmIIIA. Averages were taken over the final 300 ns of MD simulations at 298 K with disulfide bonds connected. Residues are labeled with their three-letter amino acid codes.

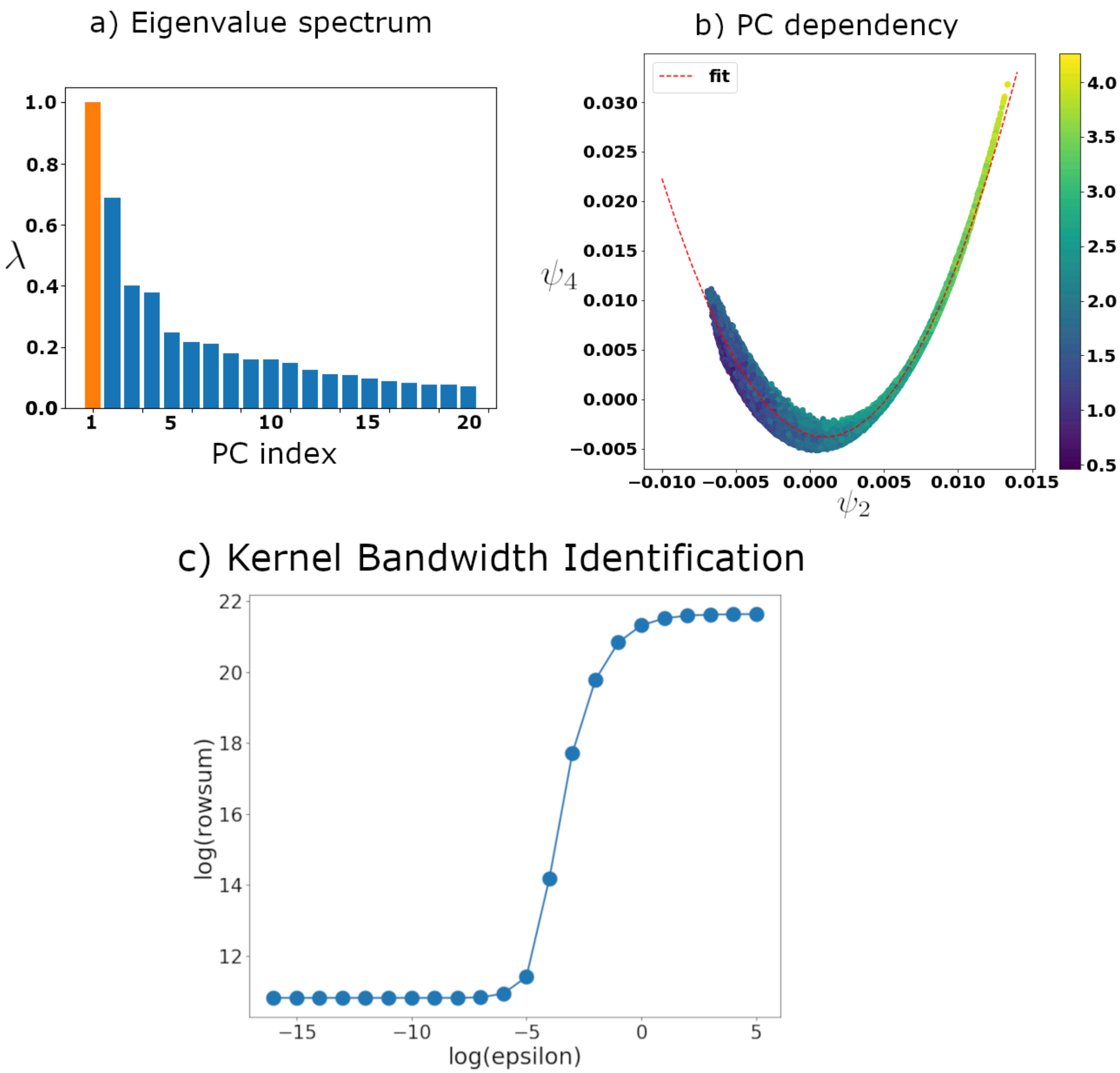


Figure S6: Eigenvalue spectrum and dependency of diffusion map performed on full set of snapshots. (a) The top twenty eigenvalues of the diffusion map matrix, with the trivial $\lambda_1 = 1$ shown in orange. We identify a gap in the eigenvalue spectrum at $\lambda_4$ by using the L method of Salvador and Chan.[80] (b) Functional dependency of two of the top three eigenvalues, along with a a polynomial fit. The arc length $\alpha_{24}$ of the curve is used to parameterize the final collective variable extracted. (c) Log-log plot of the row-sum of the adjacency matrix $\sum_j A_{ij}$ versus kernel bandwidth $\epsilon$. The best kernel bandwidth lies in the approximate linear regime and the slope of the linear regime is an approximation for the dimensionality of the system.[39]

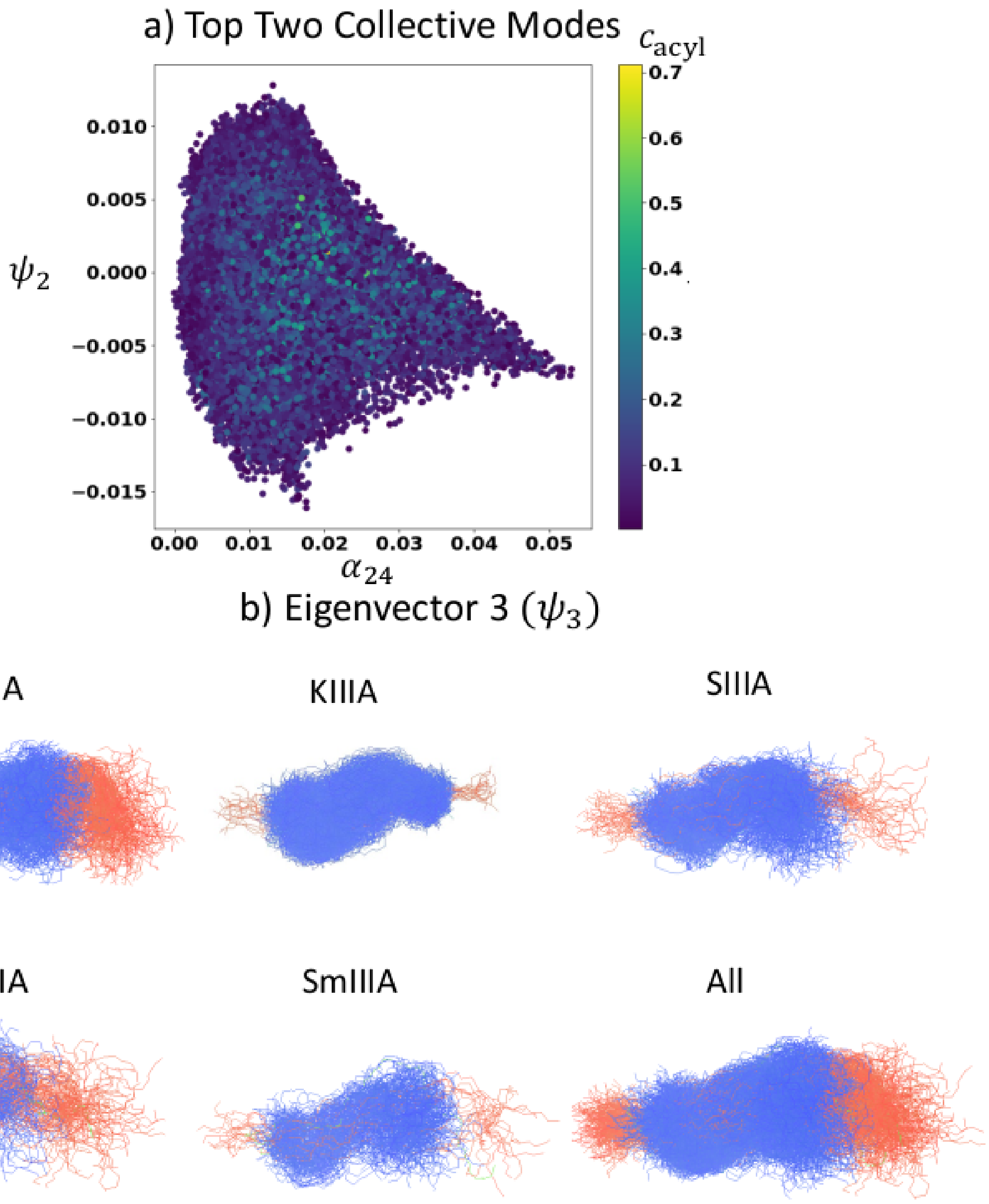


Figure S7: Collective modes identified by the composite diffusion map. In a) we illustrate the configurations projected into the two collective modes colored by the acylindricity $c_{\mathrm{acyl}}$, and in b) we supplement the visualization in Fig. 2 by displaying the configurations that contribute most strongly (top 10%) to the third top eigenvector, which is functionally dependent on the top eigenvector and correlated with the end to end length of the configuration.

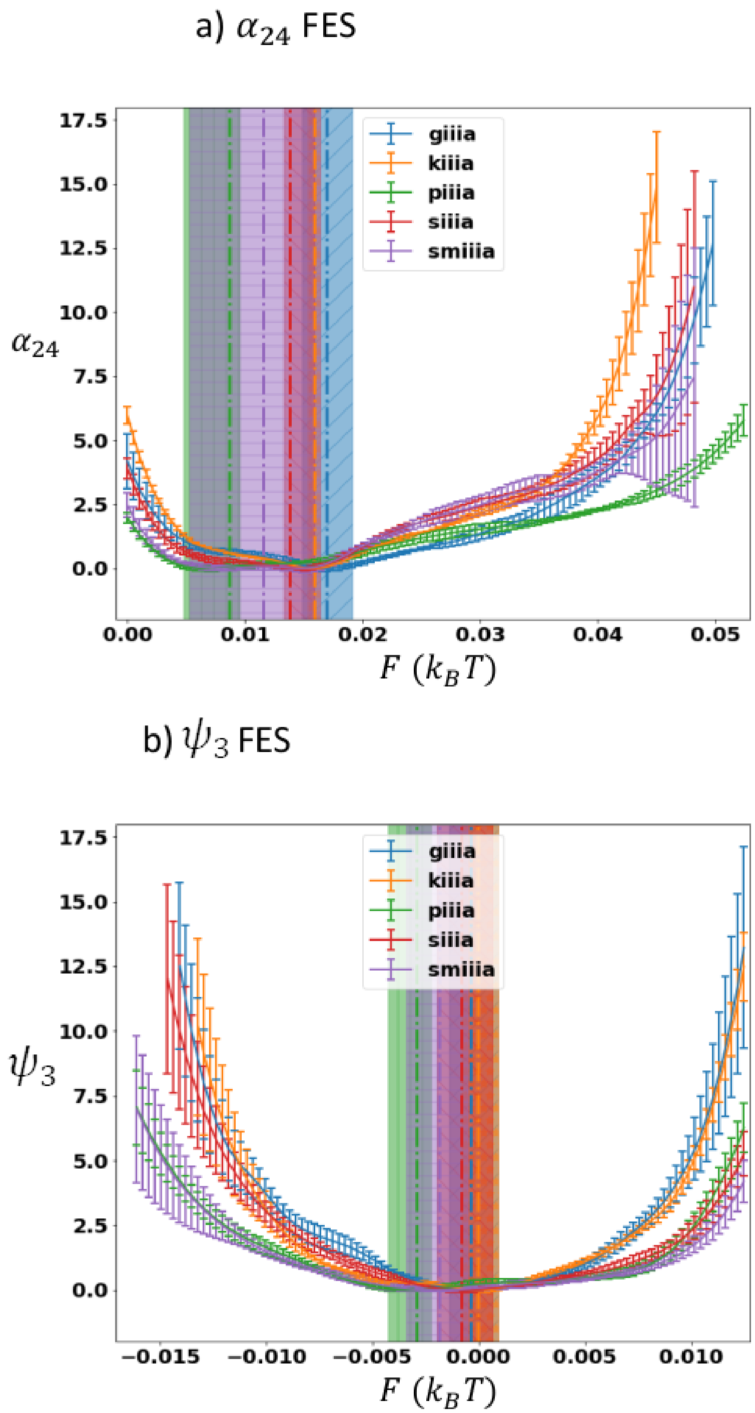


Figure S8: One dimensional free energy surfaces. In (a) and (b) we separately show the one-dimensional free energy surfaces with error bars computed by blocking with five blocks of 100 ns each. Vertical lines indicate the locations of the minima, while horizontal lines indicate the portion of the free energy surface within 5 $k_BT$ of the minima. One dimensional surfaces computed by fitting a Gaussian KDE for direct comparison between bins.

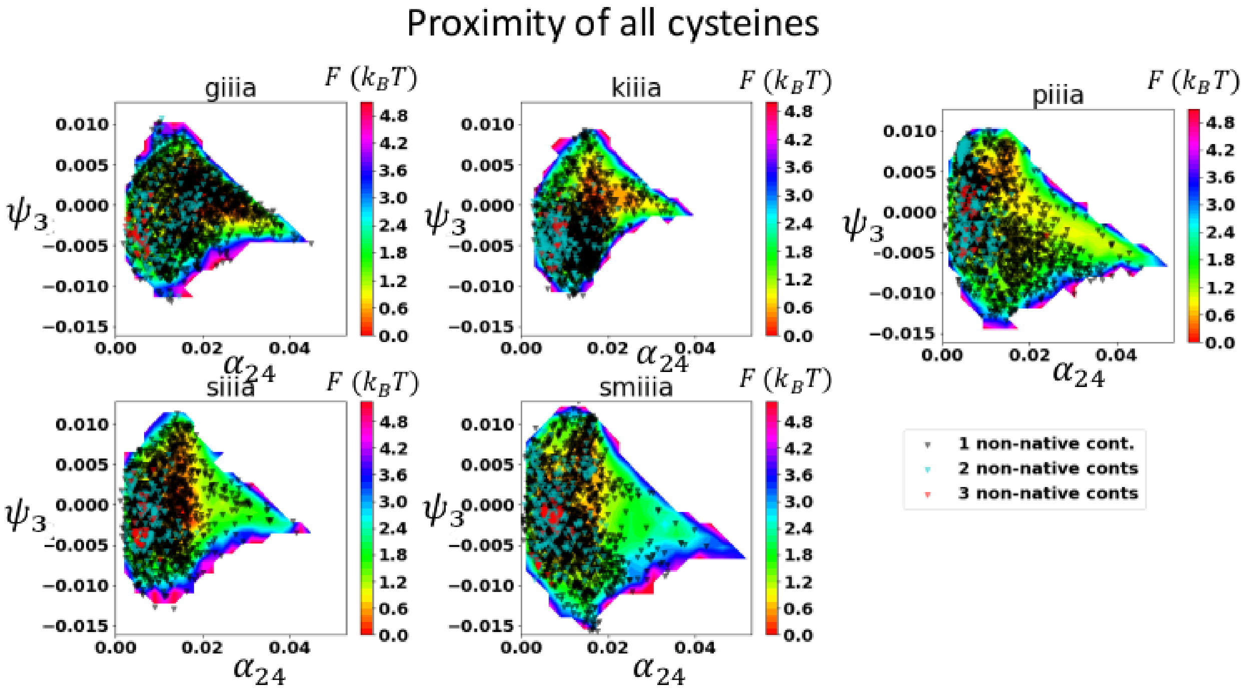


Figure S9: Differential population of transient bonds in free energy landscapes. In this figure, we indicate the portions of the free energy surfaces where cysteines are in close proximity (within a cutoff of 5 angstroms) and thus bonds might in principle be able to form between them. Black triangles indicate close proximity between one set of cysteines; cyan between two; and red between three. We indicate all proximal cysteines.

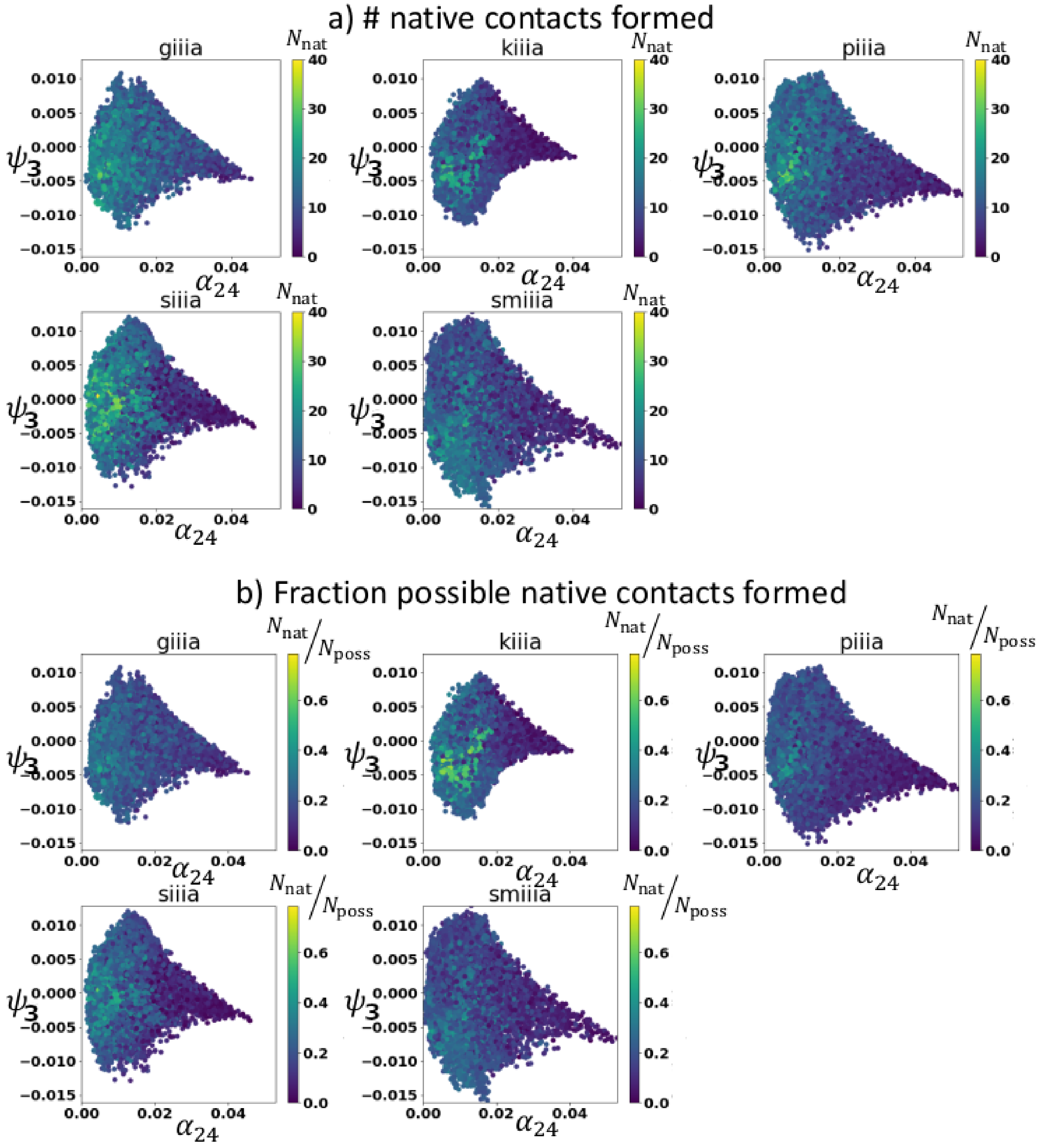


Figure S10: Native contact formation in FES. We plot the collective modes colored by (a) number of native contacts formed and (b) fraction of native contacts formed. As might be expected, a higher number of native contacts form in collapsed region of the FES (cf. Fig. 3); however, the percentage of possible native contacts formed is relatively lowless than 50% for all but KIIIA nearly across the board.

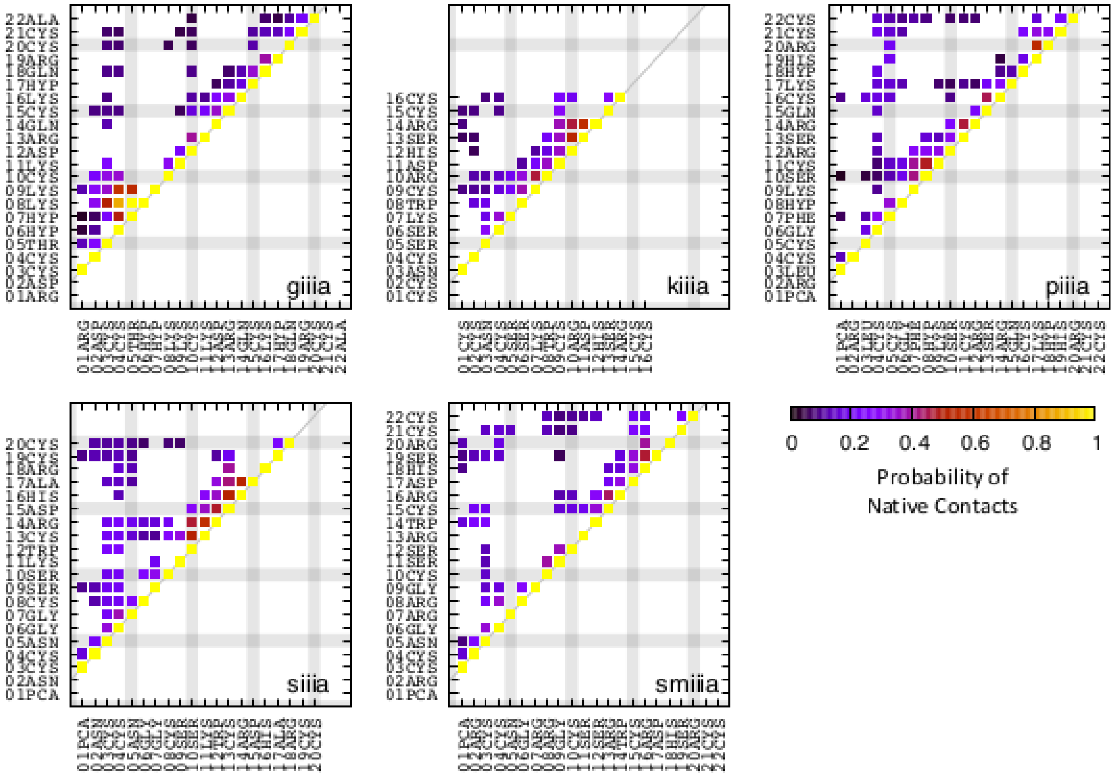


Figure S11: Probability of the native contacts for the respective conotoxins.Native contacts were identified based on the average distance between atoms from the pdb files for these conotoxins. Contact is defined as residues having heavy atoms within 6 Angstroms of one another. The probability of said native contacts was calculated via block averaging by splitting the trajectory into 10 segments. Particular structures of note include a persistent contact between THR5 and LYS8 in GIIIA, and persistent $\alpha$-helical character in residues 13-18 of SIIIA.

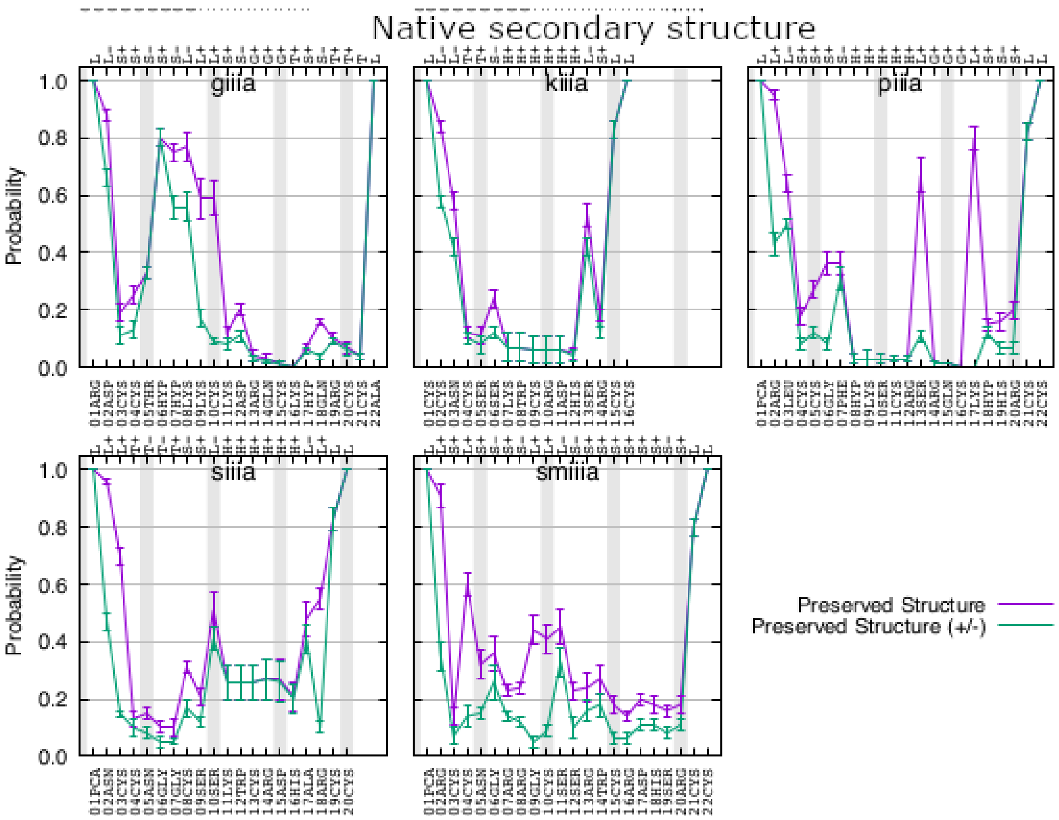


Figure S12: Probability of the secondary structure matching the observed NMR ("native") state configuration for general secondary structure (purple) and full secondary structure including handedness of the residue backbone (green).

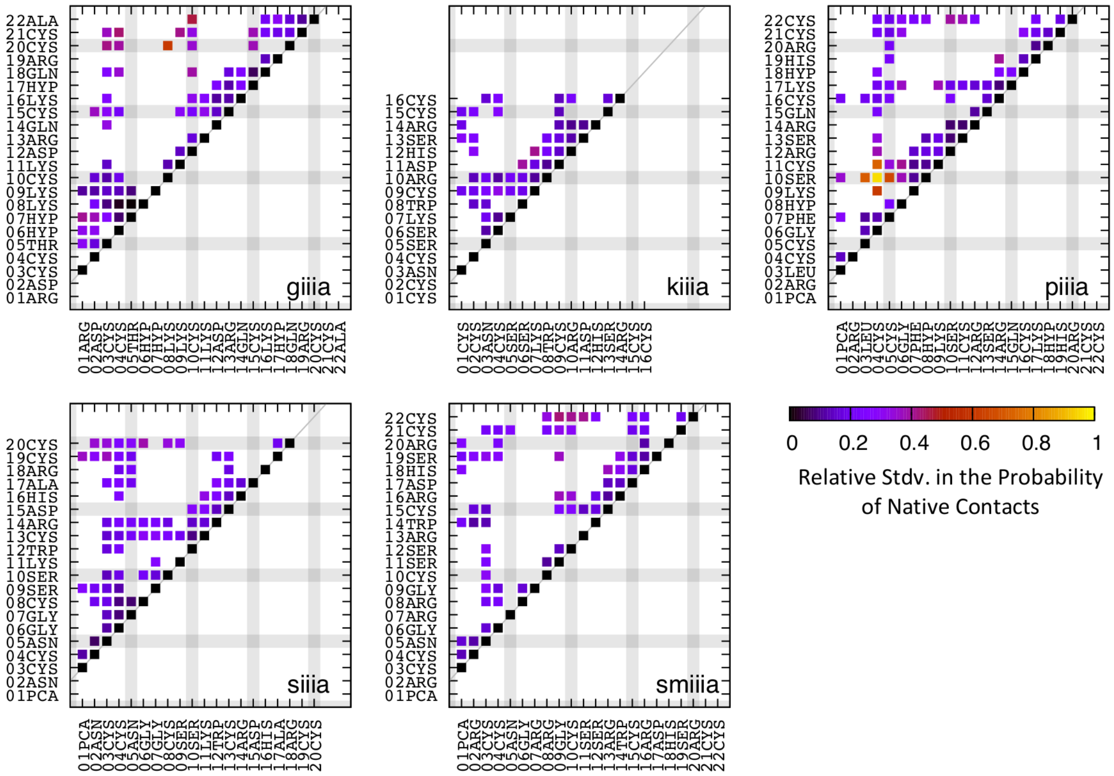


Figure S13: Relative standard deviations in the probability of the native contacts for each of the conotoxins calculated as stdv/avg.

between two atoms in the protein by extracting the principal axis of inertia $\vec{I}_3$ and finding the greatest difference $\Delta_I$ in coordinates of the atoms along that axis,

$$\mathbf{R}_3 \equiv (\mathbf{R} - \vec{r}_{\mathrm{COM}})\, \vec{I}_3 \tag{5}$$

$$\Delta_I \equiv \max(\mathbf{R}_3) - \min(\mathbf{R}_3)\,, \tag{6}$$

where $\mathbf{R}$ is the $N \times 3$ matrix of atomic coordinates, $\vec{r}_{\mathrm{COM}}$ is the position of the protein center of mass, and $\mathbf{R}_3$ is the $N \times 1$ matrix of projections along the long axis. We computed the average $\langle \Delta_I \rangle$and standard deviation $\sigma_{\Delta_I}$ of $\Delta_I$ over the final 60 ns, and set the corresponding long axis of the dodecahedral box to be $\ell_{\mathrm{long}} \equiv \langle \Delta_I \rangle + 3\sigma_{\Delta_I} + 2.4$ nm for each peptide (see Table 2 for exact values).

Table S1: Box lengths for cubic boxes for initial length characterization at high temperatures.

| **Peptide** | **Box Length** (nm) |
|---|---|
| GIIIA | 9.3 |
| GIIIA-pro | 9.3 |
| KIIIA | 8.2 |
| PIIIA | 8.2 |
| SIIIA | 7.3 |
| SmIIIA | 7.6 |
